# Functional Characterization of the Morpheus Gene Family

**DOI:** 10.1101/116087

**Authors:** Cemalettin Bekpen, Carl Baker, Michael D. Hebert, H. Bahar Sahin, Matthew E. Johnson, Arzu Celik, James C. Mullikin, NISC Comparative Sequencing Program, Evan E. Eichler

## Abstract

**DATA ACCESS:** The cDNA sequences reported in this paper have been deposited in the GenBank database (accession numbers): KF175165-KF175225 and BACs accession numbers that are used in this study: AC148621, AC190226, AC097327, AC097332, AC190226, AC097333, AC145401, AC187943, AC166855, AC166597, AC167295, AC235773, AC202644, and AC234805.

**ABSTRACT:** The burst of segmental duplications during human and great ape evolution focuses on a set of “core” duplicons encoding great-ape-specific gene families. Characterization of these gene families is complicated by their high copy number, incomplete sequence, and polymorphic nature. We investigate the structure, transcriptional diversity, and protein localization of the nuclear pore complex-interacting protein (NPIP) or *Morpheus* gene family. The corresponding core, LCRA, encodes one of the most rapidly evolving genes in the human genome; LCRA has expanded to ~20 copies from a single ancestral locus in Old World monkey and is associated with most of the recurrent chromosome 16 microdeletions implicated in autism and mental retardation. Phylogenetic analysis and cDNA sequencing suggest two distinct subfamilies or subtypes, *NPIPA* and *NPIPB.* The latter expanded recently within the great apes due to a series of structural changes within the canonical gene structure. Among Old World monkey, we observe a testis-specific pattern of expression that contrasts with the ubiquitous pattern observed among human tissues. This change in the expression profile coincides with the structural events that reshaped the structure and organization of the gene family. Most of the expressed human copies are capable of producing an open reading frame. Immunofluorescence analyses of the morpheus genes showed a primary localization to both the nucleus and its periphery. We show that morpheus genes may be upregulated upon pI:C treatment and find evidence of human autoantibodies produced against the NPIPB protein, raising the possibility that morpheus genes may be related to immune- or autoimmune-related function.

## INTRODUCTION

Gene duplication events are one of the primary sources for the emergence of novel gene functions and are, thus, fundamental to our understanding of evolution (1). The human genome shows a complex pattern of interspersed segmental duplication typified by a mosaic pattern of duplicons that arose recently from diverse regions of the genome. Approximately 430 blocks of segmental duplication are identified in the human genome (2–3). Phylogenetic reconstruction has shown that the majority of recent human intrachromosomal segmental duplication blocks have formed around a set of seven core or seed duplications (4). Notably, the core duplicons are highly transcribed and numerous human-great ape gene families have recently been described that map to these regions of the genome. These core sequences are among the most abundant and represent focal points upon which more complicated duplication segmental structures organized. Many of the genes embedded within these core duplicons show evidence of positive selection, demonstrate radical changes in their transcript expression profile compared to species with a single copy, and are highly copy number polymorphic (5–7).

The “morpheus” gene family (e.g., NPIP, NPIPL, etc.) is transcribed (5) from the core duplicon, LCR16a (low-copy repeat on chromosome 16) (8). The LCR16a core is ~20 kb in size and expanded during human-great ape lineage of evolution. In humans, it is distributed to approximately 25 distinct locations across chromosome 16 with only about one-fourth of the locations non-orthologous with chimpanzee (5). LCR16a also varies widely from 4–31 copies among great apes, which is in stark contrast to Old World monkeys where there are 1–2 copies defining the ancestral locus (5,9). Codon-based selection analysis revealed that the morpheus gene family is one of the most rapidly evolving gene families during the hominoid evolution with some exons showing as much as 10- to 12-fold excess in predicted amino acid replacement changes when compared to Old World monkey orthologs. Despite this remarkable change in protein structure, the function of this gene family is unknown. In this study, we have analyzed in detail the genomic gene/protein structure, mRNA expression, and subcellular localization and report that the morpheus gene family contains two distinct subfamilies, *NPIPA* and *B,* both of which are expressed differentially in human tissues. Interestingly, we find evidence of human autoantibodies produced against the NPIPB protein, raising the possibility that morpheus genes may be related to immune or autoimmune function.

## RESULTS

### I. Gene Structure and Protein Subtypes

We began by developing canonical gene models based on full-length transcript characterization, expressed sequence tag (EST) analysis, and cDNA subcloning/sequencing of different LCR16a loci. The analysis reveals two canonical gene structures/subtypes of the morpheus or *NPIP* gene family: Type A, termed *NPIPA,* represents the NPIP archetype (nuclear pore complex-interacting protein) as originally defined by Johnson and colleagues (5) (n = 1053 bp); while the second, *NPIPB* subtype, has a longer full-length transcript of 2544 bp (AB209632) in length (cloned in this study). The predicted molecular weight of the NPIPA protein is about 40 kDa, whereas NPIPB (e.g., NPIPL3) is 95 kDa. Although there are numerous alternative splice forms, a major difference between the NPIPA and NPIPB subtypes is the presence of exon 5 among NPIPA members, which is absent in all *NPIPB* members. In addition, *NPIPB* members show different patterns of alternative splicing leading to a 51 bp extension of exon 4, predicted to create a 17-amino acid extension sequence, as well as additional exons that become incorporated into the 5′ and 3′ ends (Figure 1). In total, the *NPIPB1* canonical gene model consists of 11 exons, of which 7 are shared with *NPIPA* members. All gene structures within the human reference genome have been renamed based on this classification according to Human Gene Nomenclature (HGNC) standards.

**Figure 1.**
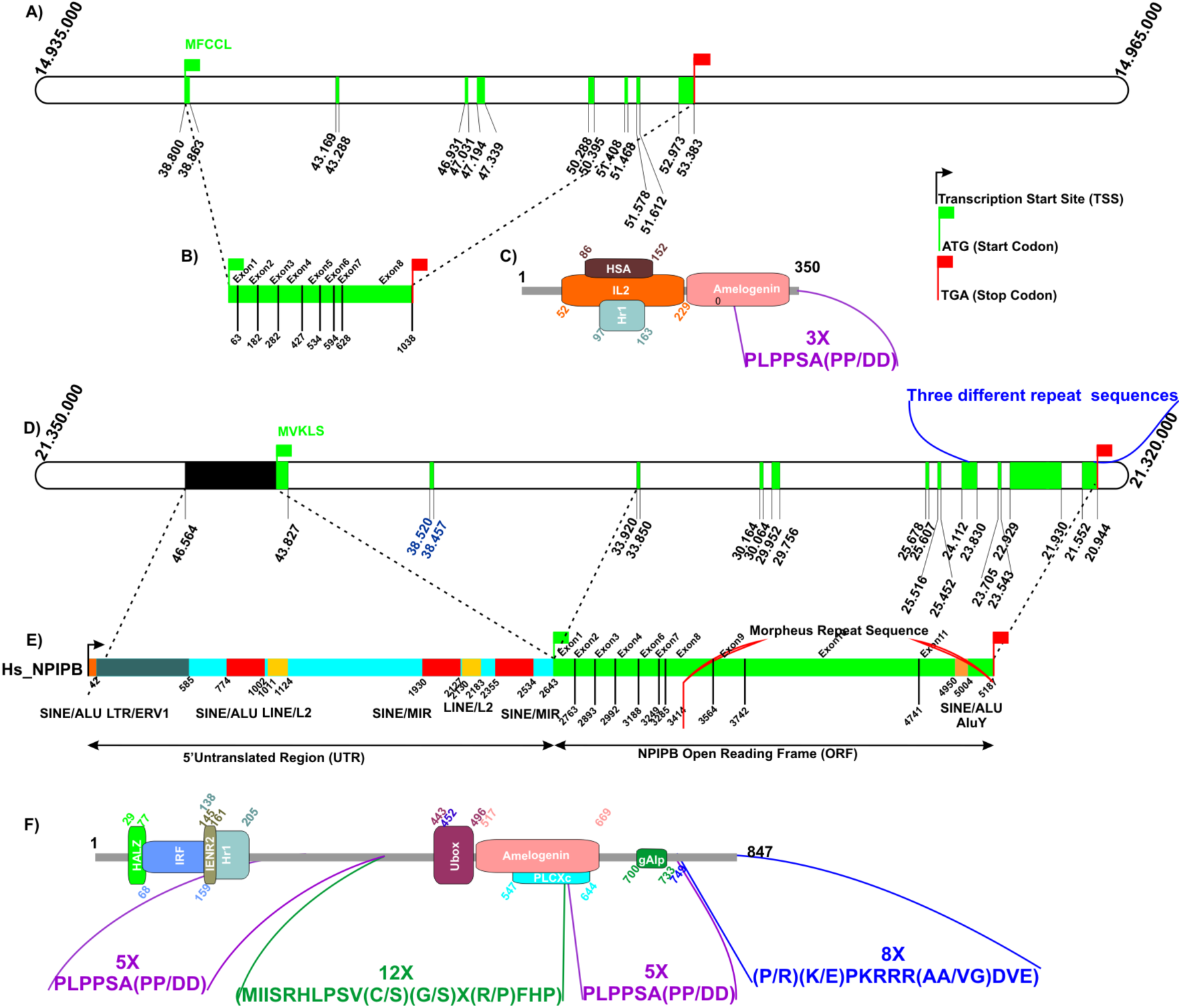
NPIPA and NPIPB gene structure. The figure shows the gene, transcript and protein structure of *NPIPA,* short, (A-C) and *NPIPB,* long, (D-F) genes. The NPIPA and NPIPB structure is based on AF132984 and AB209632, respectively. Translation start (green) and stop (red flags) codons are indicated with respect to gene structure from the human reference genome. Protein structures for the 40 kDa NPIPA (C) and 95 kDa NPIPB (F) proteins are based on SMART search database using SMART normal and genomic tools (see Supplementary Table 1 for protein domain homology). The *NPIPB* gene differs from the classical *NPIPA* gene by the presence of additional exons at the N- and C-terminal regions, eight repeat sequences, an alternative start exon at the 5′ region, and a long 5′UTR. There are, however, RT-PCR clones showing alternative splicing when compared to the classical NPIP, which is depicted in green (MFCCL). The repeat sequence composition within the 5′UTR (E) is based on RepeatMasker (Figure S3).

Conceptual translation of the two subtypes predicts very different, albeit related, protein structures (Figure 1). The beginning of the NPIPB protein consists of an alternate amino acid start sequence “MVKLS” and contains three distinct types of amino acid repeats at the carboxy (C) terminal portion of the protein: namely, two copies of PLPPSA (each 5 repetitions), 12 repetitions of a MIISRHLPSV(C/S)(G/S)X(R/P)FHP motif, and 8 repetitions of a ((P/R)(K/E)PKRRR(AA/VG)DVEP) motif. In contrast, the *NPIPA* copies carry only the hexapeptide PLPPSA repeat, which is highly variable in copy number between both NPIPA and NPIPB paralogs. In contrast, the NPIPB-specific peptides (MIISRHLPSV(C/S)(G/S)X(R/P)FHP) and ((P/R)(K/E)PKRRR(AA/VG)DVEP) appear fixed at 12 and 8 copies, respectively, among all NPIPB duplicates (Figure 1). It is interesting to note that a simple frameshift mutation (of TGAGCGTCTCTGCGGATTCCG) converts PLPPPSA to a MIISRHLPSV(C/S)(G/S)X(R/P)FHP motif, indicating that these two motifs originate from the same nucleotide sequence (Figure S1). Phylogenetic reconstruction of the C-terminal repeat sequence indicates that the mature *NPIPB* gene was derived from *NPIPA* during the process of evolution in a stepwise manner by addition of different types of repeat sequences after the split from orangutan leading to human lineage.

Protein domain analysis (as well as experimental analysis; see below) suggests that the carboxy terminus may have a role in the localization and/or association of the morpheus gene family members to the cell cytoskeleton. Recent evidence indicates that the PLPP peptide region from the Kalirin8 protein, also known as Huntingtin-associated protein-interacting protein (HAPIP), is involved in RhoGEF activity by intra and intermolecular interaction with SH3 domains (10) (Figure S2). Although the predicted protein structure of the morpheus protein is quite unique, there is 19%, 27%, 16%, and 21% similarity to the amino (N) terminal region of melanoma inhibitory activity family, member 3 (MIA3) from mouse (NM_177389), marmoset (XR_144690.1, XM_002764063), macaque (XM_001100730), and gibbon (XM_003265207), respectively. The NPIPB protein has similarity to HALZ, IRF, Hr1, to the N-terminal region of Amelogenin, and PLCXc in the C-terminal region. The shorter, NPIPA, 40 kDa protein has a significant level of similarity to HSA and IL2 in the N-terminal region (Figure 1 and Supplementary Table 1). We also predict potential phosphorylation sites, Sh3 and WW interaction sites at the C-terminal portion of the protein (Figure S2).

Interestingly, our own phylogenetic analysis using the most rapidly evolving portion of LCR16a (n = 667 bp, Figure 2) (4) confirms a remarkable structured phylogenetic relationship between and within different primate species (Figure 2). Focusing on the most rapidly evolving portion of the gene, we find that most great ape species have their own evolutionarily distinct set of morpheus genes. The NPIPB subfamily has expanded primarily within the human and chimpanzee lineages with specific branches increasing differentially. Human and chimpanzee NPIPA show a clear monophyletic origin with evidence of lineage-specific expansions in both branches. This dichotomy between NPIPA and NPIPB, however, is not predicted among the Asian apes. Only one complete copy of the *NPIPB* gene (see below) can be detected within the Asian apes and it appears to possess an incomplete gene structure; although, we cannot rule out technical problems associated with assembly of the reference genome for the Asian apes.

**Figure 2.**
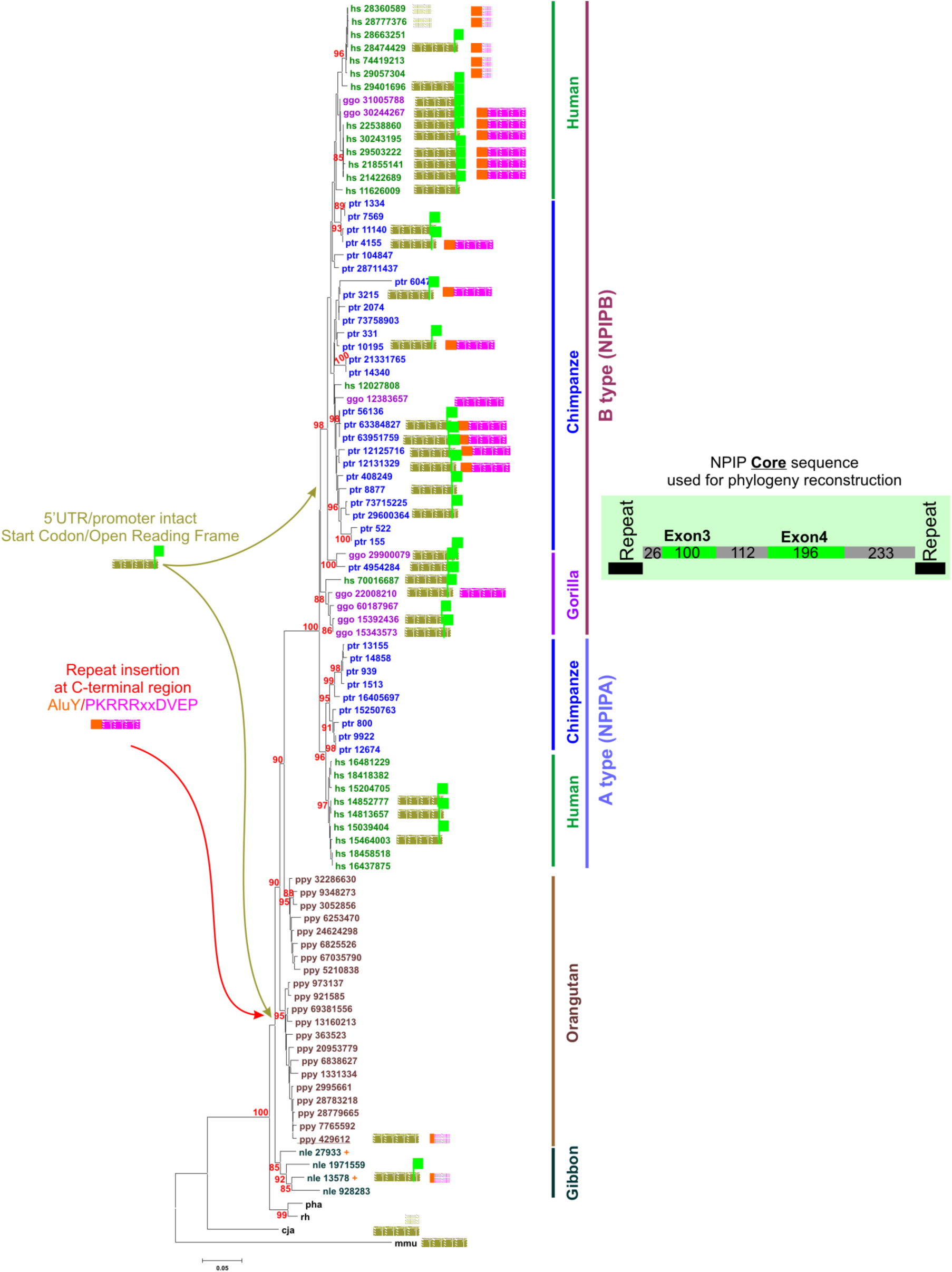
Evolutionary relationships of morpheus gene family. A neighbor-joining phylogeny constructed from a multiple sequence alignment of the most rapidly expanded portion of the LCR16a core sequence (exon-intron structure of the 737 bp fragment is shown). Species names are indicated as: mmu gray mouse lemur (*Microcebus murinus*), cja marmoset, *(Callithrix jacchus),* rh Rhesus macaque (*Macaca mulatta*), pha baboon (*Papio hamadryas*) nle Gibbon (*Nomascus leucogenys*), ppy Orangutan (*Pongo pygmaeus*), ggo Gorilla (*Gorilla gorilla*), ptr Chimpanzee (*Pan troglodytes*), and hs Human (*Homo sapiens*). Sequences were extracted from reference genome assemblies with the exception of chimpanzee. Chimpanzee NPIPB genes are mapped based in the BACs, AC148621 for 3215-R AC148538, AC148838 for 6334827-Repeat, AC097271 for 10195-Repeat AC190226, AC097270 for 4954284, AC097327 for 639517059-Repeat, AC148839 for 11140, AC097332 for 29600364, AC190226 for 12131329-R, AC097328 for 4155-Repeat, AC097333 for 12125716-Repeat and *M. murinus* NPIPB in AC235773. The number next to the respective branch name indicates the position of each core sequence within each species. Bootstrap support (n = 500) is shown only for those branches that exceed 85% and genetic distances (number of substitutions per site) were computed using the Jukes-Cantor method (20). All positions containing alignment gaps and missing data were eliminated (pairwise deletion option). Phylogenetic analyses were conducted in MEGA4 (21).

Comparative analysis of intergenic and intronic sequences indicates some radical, potentially functional, changes in the gene structure among different hominoids. We identified, for example, an Alu/ERV1 integration corresponding to the promoter region of *NPIPB* and the expansion of SINE/MIR, LINE/L2 repeats corresponding to the 5′UTR (untranslated region) of morpheus after the split of the Old World and New World monkey lineages (Figures 2, S3 and S4). Further comparative analysis of primate genomes and BAC sequence data suggested that the start codon from NPIPB emerged from noncoding DNA repetitive sequence. We find no evidence, for example, of an ATG start codon corresponding to the orthologous start position for NPIPB in either the genomes or BAC sequence from New World monkeys (*Saimiri boliviensis* (AC234805), *Callithrix jacchus* (AC202644)), Old World monkey (*M. mulatta*), or prosimian (*M. murinus*) (AC235773) (Figure S6). Additionally, no splicing acceptor region is detected in macaque for exon 2 of the *NPIPB* gene. Combined, these data suggest that the *NPIPB* model emerged specifically within the ape lineage. We identified and sequenced multiple BACs carrying copies of morpheus from gibbon (*N. leucogenys*) and orangutan (*P. pygmaeus*). Four gibbon BACs (AC187943, AC166855, AC166597 and AC167295) showed evidence of a repeat-rich 5′UTR and ATG start codon associated with the *NPIPB* model. Only one BAC (AC145401) in orangutan (mapping to ponABe2 chromosome 16 position 429612) contains the *NPIPB* 5′UTR (Figures 2 and S5) but this single *NPIPB* paralog does not possess a start codon consistent with canonical *NPIPB* model, suggesting that it is not capable of producing a functional *NPIPB* gene product. It is possible that this copy represents the ancestral locus from which African great ape copies of NPIP emerged.

### II. Transcript Expression Analyses

Based on these morpheus gene models, we performed comparative RT-PCR from cDNA derived from various macaque and human tissue material. Macaque expression is predominantly testis (as well as weak thymus expression) in contrast to human, which shows a strong ubiquitous pattern of expression (Figure 3). We quantified mRNA expression levels in human tissues by real-time quantitative PCR (RT-QPCR) and observed similar levels among the tissues (2- to 7-fold). Liver and heart showed the lowest, while thymus, testis and brain showed the highest levels of expression (Figure 3). We cloned and sequenced a total of 56 RT-PCR products from HeLa and A549 cells and, based on unique sequence differences, assigned them to specific copies on human chromosome 16 (Figure 4). We find that at least 13 of the 24 family members are transcribed revealing a remarkably complex pattern of 5′ alternative splice forms (Figures 4 and S9). Combined with EST analysis, there is evidence for at least five alternative start exons for this gene family. We predict that some of these alternate first exons lead to a premature stop codon and nonfunctional transcript. For example, the PKD1-NPIP fusion transcripts are predicted to lead to a pseudogene as a result of a splicing frameshift. In total, about 80% of sequenced cDNA maintain an open reading frame consistent with the protein models described above while 20% (11/56 clones) show evidence of a frameshift. Although it is not always possible to unambiguously assign each transcript to a specific locus, there is evidence that some loci are producing both functional and nonfunctional transcripts due to alternative splicing (Figure 4).

**Figure 3.**
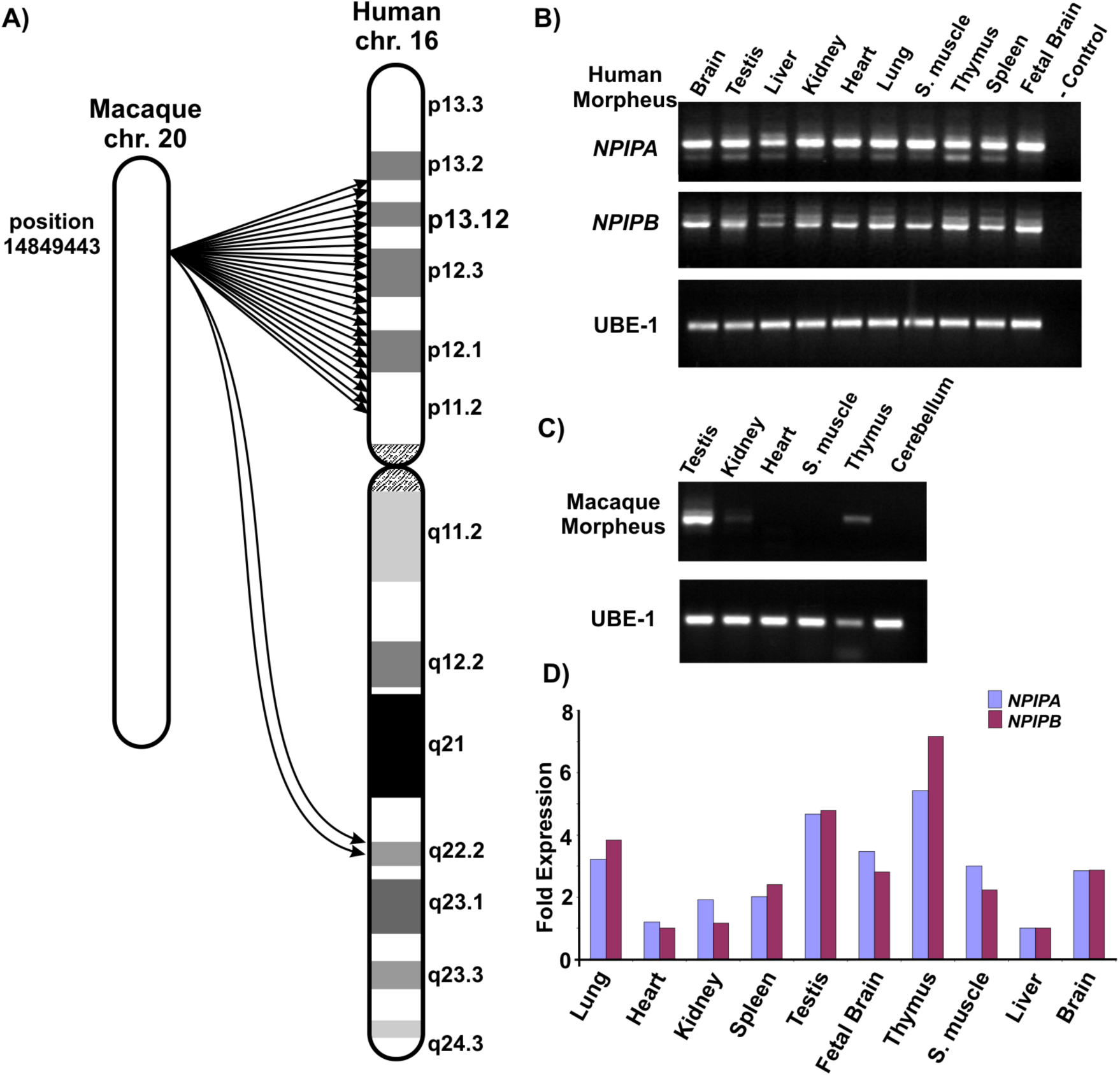
Expression analysis. (A) The location of morpheus gene duplications are depicted in the context of an ideogram for rhesus macaque (left) and human (right) based on macaque and human genome reference sequence (rheMac2 and GRCh37, respectively). (B) NPIP RT-PCR results from cDNA prepared from total RNA extracted from various human tissues are compared to (C) rhesus macaque tissues. RT-PCR was compared to *UBE1* expression as a positive control. (D) Real-time PCR analysis of morpheus gene family. Quantitative PCR analysis to show differential expression of morpheus gene family within human tissues is presented. Liver is used for normalization, which is the lowest *NPIP* expressing tissue. *UBE1* is used for control. There is no significant enrichment of either long, (NPIPB), (95 kDa) or short, (NPIPA), (40 kDa) version of the morpheus mRNA copy present in any tissue tested. Morpheus is expressed relatively high level (approx. twofold) in testis and thymus both in human and macaque. The primers for real-time PCR were designed specifically for exon 6 and 8 NPIP_Exon6F, and NPIP_Exon8R (Supplementary Table 2). To distinguish both types of morpheus genes, the specific primer set for each copy is designed based on reference assembly.

**Figure 4.**
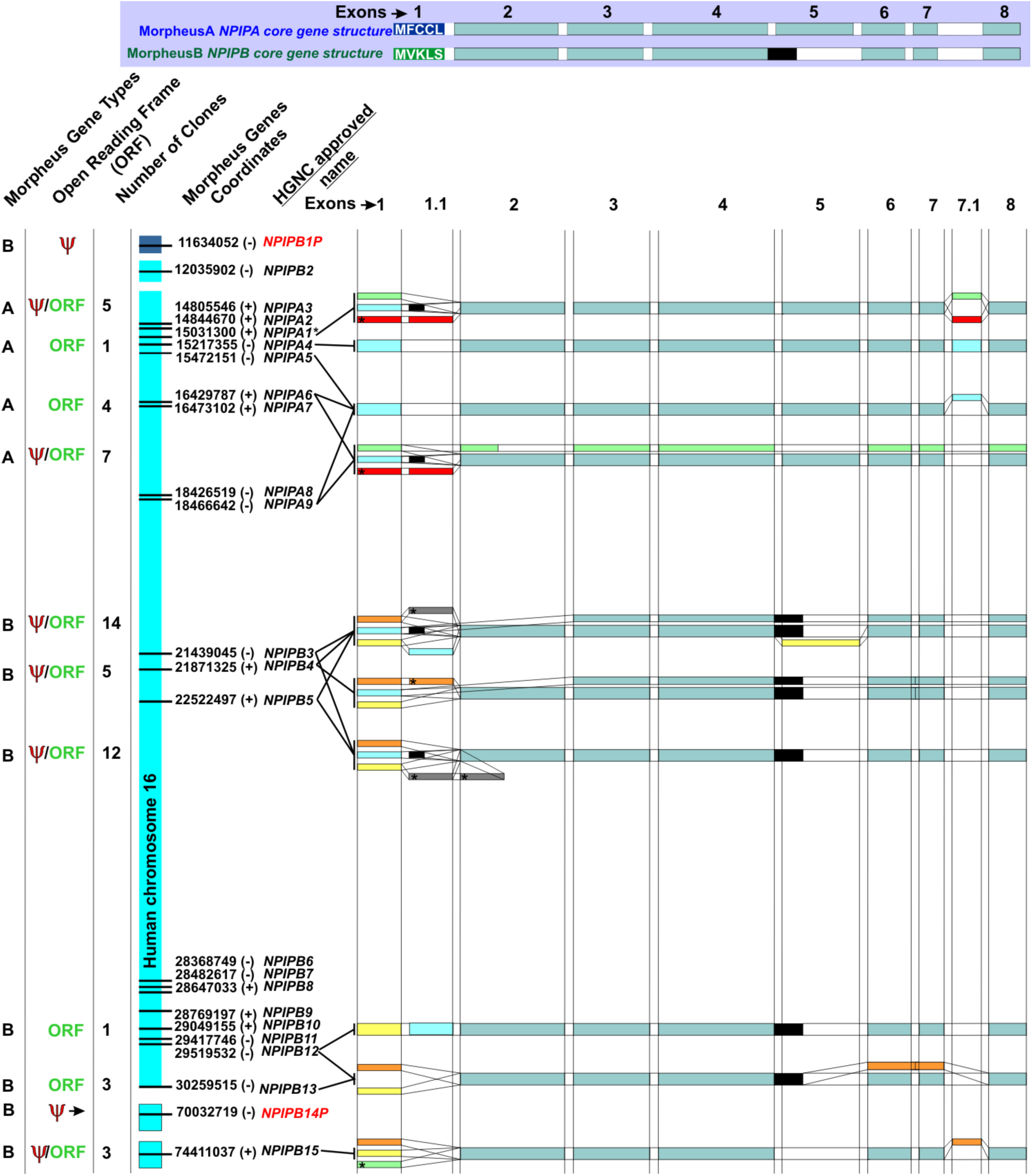
High-resolution mapping of the morpheus gene family on a human chromosome. Mapping of the individual morpheus gene family members based on comparison of cDNA clone and human genome sequence. All genes and pseudogenes are assigned to either NPIPB or NPIPA subfamilies based on differences in the gene structure and named according to HGNC standards. Primers were designed for exon 1 and the beginning of exon 8 (before the repeat region of morpheus) and a total of 55 RT-PCR products were cloned and sequenced. The exon 1 primers were designed to amplify both canonical and alternative first exons as detected by 5′RACE and EST analysis (Supplementary Table 2). Transcripts corresponding to a likely pseudogene due to an incomplete open reading frame are shown in red. The position of each gene is depicted with a horizontal line and the (+/-) sign indicating the orientation of transcription (5′ to 3′) with respect to human chromosome 16. Clones that cannot be assigned to a unique genomic location are indicated by multiple arrows.

Because of the higher levels of expression in the thymus and because immune response genes often show signatures of adaptive evolution (11–14), we investigated whether morpheus transcripts could be differentially regulated in various immune response pathways. We tested this by assessing morpheus total RNA expression levels of HeLa cell lines before and after stimulation with immunogenic agents. This included cytokines, such as interferons (α/β and γ), TNFα, and PAMPs (e.g., lypopolysaccaride (LPS), a mimic of pathogenic bacteria, and polyinosinic-cytidylic acid (pI:C), a mimic of viral RNA). We designed specific primer sets to assess three distinct regions of the morpheus gene model in order to span most of the major splice forms. Upon transfection of HeLa cells with pI:C (1μg/ml) for 6 hours, we observed a reproducible upregulation of morpheus mRNA levels of about sixfold (Figure S10). None of the other immunogenic stimulant agents induced higher levels of mRNA (data not shown).

### III. Protein Analyses and Subcellular Localization

We investigated the subcellular localization of the protein by generating a series of rabbit polyclonal antibodies against one peptide that is specifically recognizing NPIPB; MP2:CSPEPKRRRVGDVEP, one peptide specific for NPIPA; N5:HGEKERQVSEAEENGK, and two peptides recognizing all types of morpheus: MP4:CTKVNRHDKINGKRKT, N8:CRMAAVQHRHSSGLPYW, respectively. We show that both N8 and N5 antibodies can specifically immunoprecipitate recombinant the GFP-NPIP fusion protein from HeLa cell lysate (Figure S11). The specificity of the N8 antibody recognition is also confirmed by immunofluorescence analysis of transiently transfected HeLa cells with the GFP-NPIP fusion gene construct. Similarly, the specificity of MP4 and MP2 is determined by immunofluorescence analysis of overexpressed *NPIPB* flag-tagged at the N-terminus in HeLa cells (data not shown). The endogenous protein analysis using the N8 antibody gives specific band recognition at 95 kDa and 43 kDa in human tissue lysate when compared to mouse testis tissues (Bl6) (Figure S12).

Analysis of the putative protein domain architecture predicts at least four nuclear localization signals (NLS) present in the middle and at C-terminal region of the NPIPB protein (Figure S13). This is in contrast to NPIPA where there is no sequence similarity to any known NLS. To test the NLS prediction for the NPIPB protein, we successfully cloned partial copies of the two *NPIPB* genes (K1.3 (maps to chr 16:28671295) and K1.4 (maps to chr 16:22432624)), which differ only in exon 2 and the C-terminal region (Figures S14 and S15), into a mammalian expression vector (pM1-MT Roche) as well as the shared (seven-exon) region between NPIPA and NPIPB (c12 construct).

Despite multiple attempts, we were unable to clone a full-length copy of NPIP that included the entire C-terminal repeat region of the gene model. Transient overexpression of NPIPA constructs, whether tagged N-terminally with GFP or flag, were primarily localized to the nucleus in HeLa cells (Figure 5). There is a partial localization around the periphery of the nucleus but the majority of the signal localizes within the nucleus. In contrast, the K1.3 clone of *NPIPB* localizes exclusively to the nucleus while the K1.4 construct, corresponding to the other *NPIPB* subtype, shows signal primarily in the cytoplasm (Figure 5). Notably, the shared *NPIPA/NPIPB* gene model construct, c12, clearly aggregates around the nuclear periphery (Figure S16). The major difference between *NPIPB* and c12 is the presence of additional 5′ and 3′ exons (Figure 1), which are not present in c12 (Figure S14). c12 also differs from GFP Flag-NPIP by lacking the NPIP-specific exon 5 and the repeat sequence at the C-terminal region (Figure S14).

**Figure 5.**
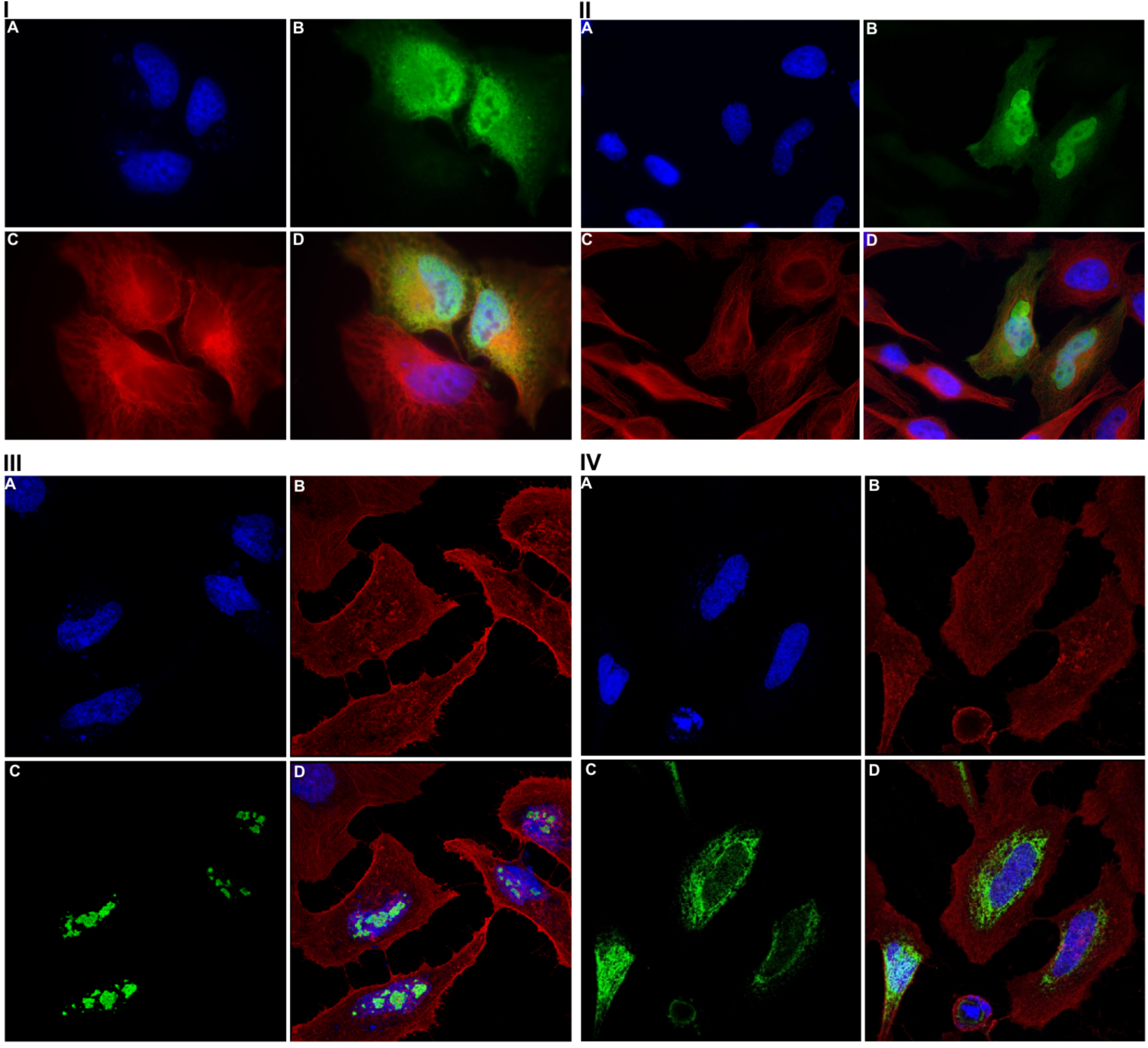
Immunofluorescence analysis of NPIP constructs in HeLa cells. I) (A) HeLa cells are transiently transfected using the FuGENE HD transfection reagent (Roche) with GFP-NPIP for 24h and stained with a 1–100 dilution of N8 antibody (B) and a 1–1000 dilution of anti-tubulin antibody. (C) Primary antibody staining is performed overnight at 4°C. Nucleus (A) is visualized by DAPI staining. Merged figure (D) shows the co-localization between N8 (B) and tubulin antibody (C) staining. The majority of the HeLa cells show specific staining within the nucleus. However, a few cells show accumulation of transfected morpheus protein around the nucleus. II) Immunofluorescence analysis of Flag-NPIP fusion gene in HeLa cells. HeLa cells are transiently transfected by using the FuGENE HD transfection reagent (Roche) with Flag-NPIP for 24h and stained with 1–100 dilution of N8 antibody (B) and 1–1000 dilution of anti-tubulin antibody (Sigma) (C). Nucleus (A) is shown by DAPI staining. Merged figure (D) shows the co-localization between N8 (B) and tubulin antibody (C) staining. The majority of the HeLa cells are showing specific staining within the nucleus. III) Immunofluorescence analysis of morpheus *B* (K1-3) in HeLa cells. HeLa cells are transiently transfected by using the FuGENE HD transfection reagent (Roche) with K1-3 for 24h and stained with 1–500 dilution of MP4 antibody (C) and 1–1000 dilution of anti-beta-actin antibody (Cell Signaling) (B). Nucleus (A) is visualized by DAPI staining. Merged figure (D) shows the co-localization between MP4 (C) and beta actin antibody (B) staining. The majority of the HeLa cells are showing specific staining within the nucleus. IV) Immunofluorescence analysis of *NPIPB* (K1-4) in HeLa cells. HeLa cells are transiently transfected by using the FuGENE HD transfection reagent (Roche) with K1-4 for 24h and stained with 1–500 dilution of MP4 antibody (C) and 11000 dilution of anti-beta-actin antibody (mouse monoclonal, Cell Signaling) (B). Primary antibody staining is performed for overnight at 4°C. Nucleus (A) is visualized by DAPI staining. Merged figure (D) shows the co-localization between MP4 (C) and beta actin antibody (B) staining. See (Figure S14) for the details of the morpheus constructs.

Combined, these findings suggest that the C-terminal portion of NPIPB is required for the proper localization of the morpheus protein to the nucleus. Although, no NLS was predicted for NPIPA, exon 5 may contain signals for proper localization of the NPIPA protein around the nucleus. Additionally, we have observed a proportion of the cells having cytosolic localization when the K1-4 construct is overexpressed. The unpredicted subcellular localization of the *NPIPB* type K1-4 construct shows that the presence of exon 2 and the C-terminal repeat sequence may be important in this regard, because it is the only place where the two constructs differ (Figures S14 and S15). It could also be possible that single amino acid substitutions observed within K1-3 and K14 may be responsible but this needs to be confirmed by further functional analysis.

The immunofluorescence analysis above was also confirmed by western blot analysis (Figure 6). We find that the 95 kDa band is detected only in nuclear/cytoskeleton/insoluble fraction consistent with the protein domain analysis showing that the N-terminal portion of the 95 kDa NPIPB protein has a domain structure similar to HALZ and IRF transcription factors. In addition, immunohistochemistry analyses using MP2 antibodies show that the endogenous NPIPB protein localized primarily within the nucleus in HeLa cells (Figures 5, 6 and S17; see below). In a recent screen for B cell lymphoma-associated antigens, Cha and colleagues, unknowingly cloned four different constructs corresponding to a specific auto-antibody fraction of the isolated IgG purified from both treated and normal individuals (15). Our detailed analysis shows that, in fact, clones FL-aa-4-1, FL-aa-4-2, FL-aa-4-3, and FL-aa-4-4 correspond to the C-terminal repeat region of the NPIPB protein. To confirm these findings, we transiently overexpressed the morpheus gene construct K1-3 in HeLa cells and co-stained transfected constructs with MP2 and MP4 antibody together with an isolated IgG fraction from a healthy donor. We found that the isolated human IgG from a healthy adult donor can specifically recognize *NPIPB* genes K1-3 (Figure S18).

**Figure 6.**
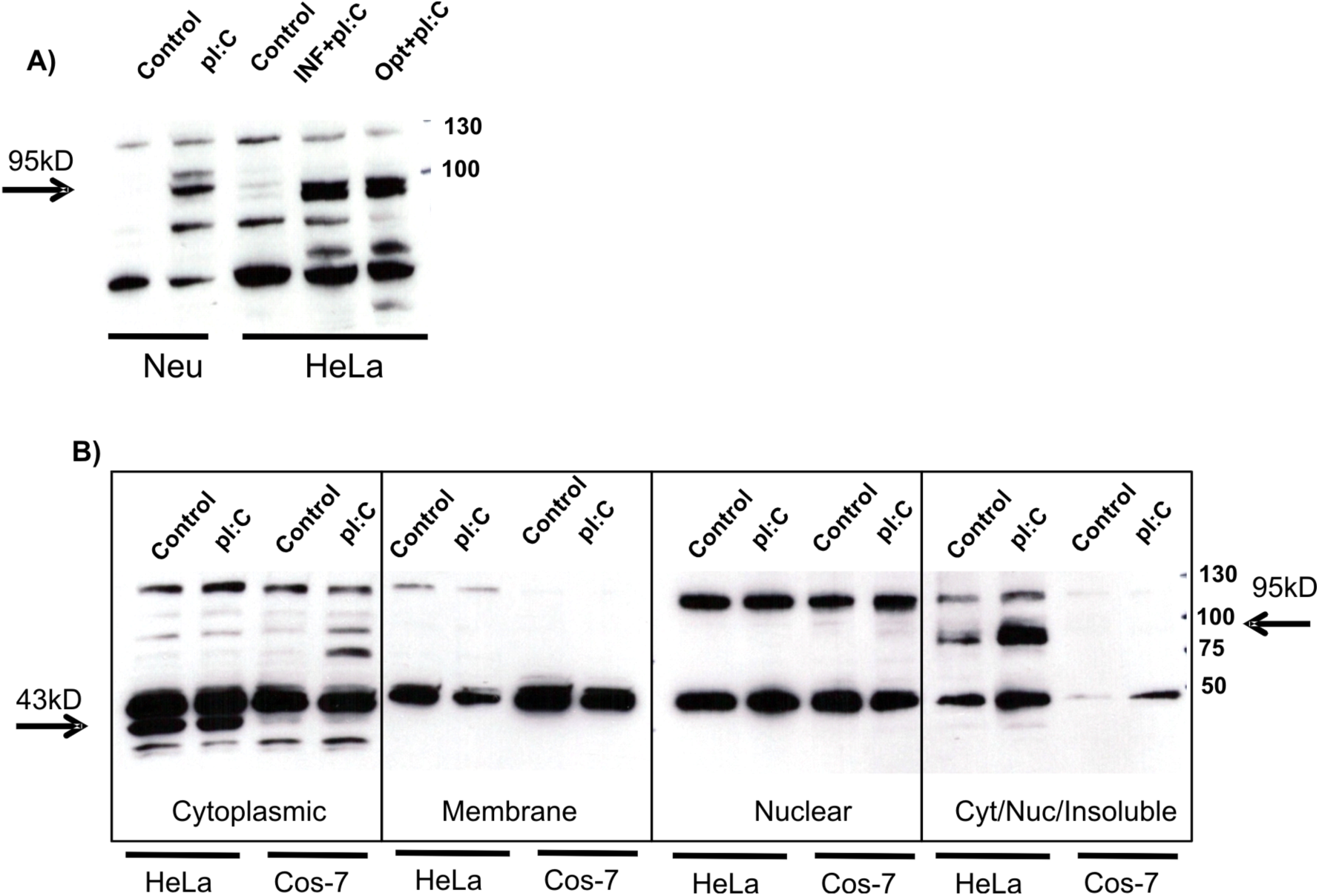
Subcellular fractionation of cell lysate treated with pI:C treatment for 24h. Subcellular fractionation of HeLa cell lysate was separated by SDS-PAGE and blotted on a nitrocellulose membrane. A 1 to 500 dilution of N8 antibody is used to detect specific signal on a western blot. Arrows indicate the position of bands corresponding to 43 kDa and 95 kDa after pI:C treatment of Neuroblastoma (Neu) and HeLa cells incubated with pI:C, pre-incubated for 6 hours with interferon alpha (INF+ pI:C) and Optimum Glutamax media (Invitrogen) before treatment of pI:C for 24h. (A) In the case of HeLa and neuroblastoma cells, only the nuclear cytoskeleton fraction was used for the western blot analysis. (B) Subcellular fractionation of HeLa and Cos-7 cells. Cos7 cells that do not contain morpheus are used as control for pI:C treatment. In Cos7 cells, no morpheus staining is observed around 95 kDa in the fraction of Cyt/Nuc/Insoluble (right).

### IV. Stable Transfection Analysis

To characterize functional properties of morpheus proteins *in vivo,* and toxicity of overexpression of morpheus, we generated stable cell lines expressing the recombinant morpheus protein in mouse embryonic fibroblast (MEF) cells under the control of a Tet-Off inducible gene expression system. In the Tet-Off inducible gene expression system, cells were grown in the presence of tetracycline homolog doxycyline, and upon depletion of doxycyline, synthesis of recombinant morpheus protein is induced. Immunohistochemistry analyses of MEF cells that are stably expressing the morpheus gene (*NPIP*) under the control of Tet-Off promoter system show no or trace amounts of protein accumulation in doxycyline depleted cells for 24 hours unless the cells are transfected with pI:C using the Lipofectamine^®^ 2000 transfection system (Invitrogen). This suggests that the recombinant morpheus protein might be degraded immediately in mouse cells. Upon transfection of NPIP-MEF cells with pI:C (1 μg/ml) for 6 hours, the morpheus protein is detected in the periphery of the nucleus and nuclear membrane similar to the results reported by Johnson and colleagues (5) (Figure 7). The total ratio for protein accumulation after transfection within MEF cell population was approximately 10–15%, potentially linked to transfection efficiency. We repeated the experiment using another morpheus-specific antibody (N5) and unrelated gene construct (luciferase gene) with similar results under the same conditions (data not shown).

**Figure 7.**
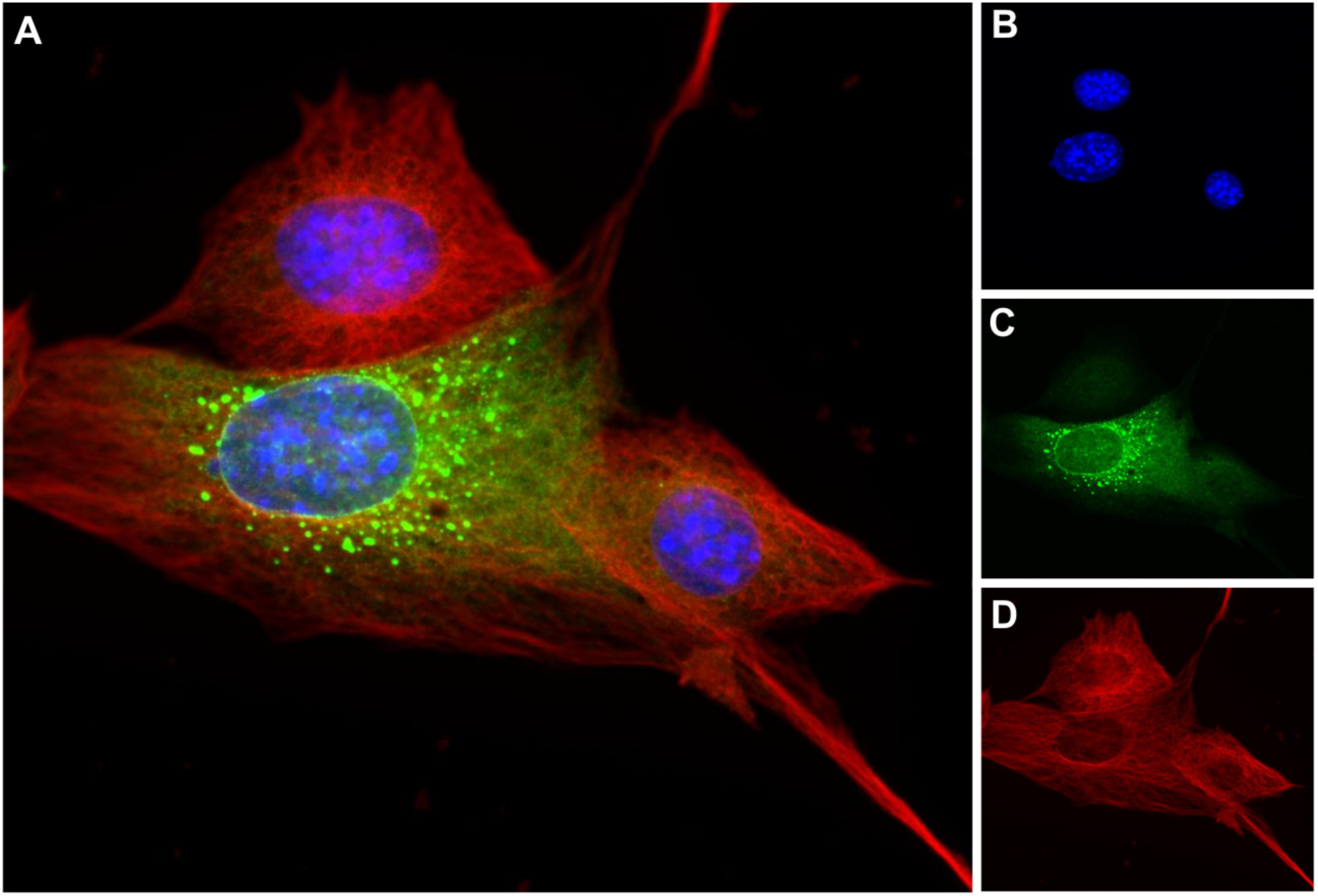
Immunofluorescence analysis of pI:C transfected stable Tet-Off NPIP-MEF cells. Immunofluorescence analysis of the morpheus gene in mouse embryonic fibroblast (MEF) cells stably expressing NPIP under the control of Tet-Off inducible promoter. NPIP-MEF cells are grown in the absence of doxycycline, G418 and hygromycin for 24h before transfection. Six hours after transfection of NPIP-MEF cells with pI:C (1 μg/ml), cells were fixed with cold (20°C) MeOH for 5 min. The merged figure (A) shows the pI:C transfected Tet-Off inducible NPIP-MEF cells stained with morpheus-specific N8 antibody (green). The nucleus is stained with DAPI (blue) (B), polyclonal antibody (N8) against morpheus (green) (C); polyclonal antibody against α-tubulin (red) (D) is used as positive control.

## DISCUSSION

The morpheus gene family is one of the most rapidly evolving gene families to have emerged during human-great ape evolution (5). In this study, we attempted to systematically characterize differences in expression, structure, and subcellular localization for major isoforms in order to gain insight into its evolution and potential function. Cloning and sequencing of RT-PCR products of the morpheus family showed that at least 13 of 25 copies in the human genome are actively transcribed in cultured human cell lines (HeLa and A549 cells). Our gene structure analysis suggests that members of the gene family may be operationally divided into two distinct subgroups: *NPIPA* and *NPIPB.*

The NPIPA subfamily, as originally reported by (5), is simpler in structure consisting of eight exons, including a diagnostic in-frame exon, “exon 5”. Members of the second group, NPIPB (or morpheusB), are typically larger (10 exons), are transcribed by an alternate promoter, and possess a repeat-rich 5′UTR as well as additional 3′ exons, including three distinct repeat sequences at the C-terminal portion of the gene. Comparative sequence analyses predict an expansion of the NPIPB subtype within great apes after the split from the orangutan lineage. This rapid expansion of the *NPIPB* gene coincided with the emergence of a start codon (first exon) and numerous retrotransposition events within the 5′ and 3′ ends of the gene (Figures 2 and S5). Since no functional *NPIPB* gene is detected within the New World or Old World monkey genomes, we suggest that the NPIPB subtype derived from duplication of NPIPA followed by the accumulation of numerous structural and single-basepair variants. In particular, the insertion of an AluY element near the C-terminus followed by a frameshift mutation led to the evolution of an entirely new amino acid repeat motif, PKRRRxxDVEP, associated specifically with the NPIPB subfamily.

Transcripts from both morpheus subtypes are expressed widely in all human tissues in contrast to other primates (e.g., macaque), which show a much more restricted pattern of expression (Figure 3). If the macaque expression profile is considered ancestral, our analysis suggests that the morpheus gene family evolved from testis- and thymus-specific mRNA expression to a ubiquitous pattern in concert with its expansion in copy number. It is possible that the accumulation of common repeats near the 5′UTR and/or the juxtaposition of novel segmental duplications facilitated this broader expression profile in the ape lineage as has been observed for other gene families (16–17). Despite its broad expression profile in humans, Homan and colleagues identified *NPIP* as one of two gene families highly expressed in the macula of the retina (18) where its transcriptome abundance in photoreceptors is second only to that of the opsin genes. They suggested the gene family as a potential candidate for age-related macular degeneration.

The NPIPA protein was originally reported to localize to nuclear membranes where it potentially interacted with nuclear pore proteins (5). In this study, we generated polyclonal antibodies against peptide sequences specific to both NPIPA and B subtypes. Predicted protein domain analyses, subcellular localization experiments, and overexpression studies confirm a translated product that primarily localizes to the nucleus or its periphery. Interestingly, actively transcribed members of the NPIPB subfamily localize predominantly to the nucleus in human cells and we show that the C-terminal region of the NPIPB protein is important for this nuclear localization. It is possible that these rapid changes in the morpheus protein structure evolved to direct subcellular trafficking between the cytosol and the nucleus.

Surprisingly, we also find that healthy humans produce an auto-antibody against the C-terminal of NPIPB proteins (Figure S18), confirming previous findings although the authors did not associate the target with NPIP (15). Recognition of self-antigens in human autoimmune diseases involves both T and B cell responses and may suggest a role for *NPIP* genes in immunity or autoimmunity. In this light, it is noteworthy that when stably transfected NPIP-MEF cells are exposed to pI:C—a known immunostimulant for TLR3—the morpheus protein localizes to a vesicular structure and accumulates around the nuclear membrane. In addition, the N-terminal region of the long (95 kDa) morpheus protein has significant similarity to a transcription factor binding protein HALZ, which is a homeobox-associated protein, and IRF7, which is an interferon regulatory protein (Figure 1 and Supplementary Table 1). One possibility may be that the morpheus protein is activated by the pI:C-dependent immune response pathway. Upon activation, the morpheus protein is modified (phosphorylated or glycosylated) and transported to the nucleus where it is predicted to maintain its function. The morpheus protein may first need to be stabilized by post-translational modification and later transported to the nucleus with the help of other, as of yet unknown, human-specific factors.

## MATERIAL AND METHODS

### RT-PCR

cDNA was prepared using the RT-PCR kit (Roche) according to the manufacturer instructions. Total RNA used for cDNA preparation was extracted from Rh (macaque) tissues, HeLa and A549 cells (RNA easy, Qiagen), and the total RNA for hs (human) tissues (Clontech). RT-PCR products prepared using the cDNA from the cells were cloned to the pGEM-T-Easy vector. Colony PCR was used to identify positive clones. These were selected for mini-prep plasmid isolation and sequencing. In total, 96 colonies from A549 and 81 clones from HeLa cells were analyzed. UBE1 are used as positive controls. PCR was performed in 20 μ1 reactions composed of 0.8 μl of a 10 μΜ dilution of the forward primer and reverse primer, 10 μl of Roche (11636103001) PCR Master Mix. The following PCR conditions were used: 1 min at 94°C, followed by 38 cycles at 94°C for 30 sec, 55°C 30 sec, and 72°C for 30 sec followed by 7 min at 72°C.

### Real-time PCR

Morpheus splice variants were detected by a quantitative PCR assay using the LightCycler SYBR Green System (Roche) with exon 6 and 8 primers (see Supplemental Table 2 for a complete list of oligonucleotides used). cDNA was synthesized using mRNA prepared from lymphoblast cell lines. The amount of measured transcripts was normalized to the amount of the GAPDH and UBE1 transcript. The following real-time PCR conditions (C) were used: 3 min at 95°C, followed by 50 cycles at 95°C for 15 sec, 55°C 20 sec, and 72°C for 20 sec.

### Stable cell line preparation

We generated an inducible Tet-Off cell line stably expressing morpheus C5 (Classical NPIP), C3 and C4 (mutated versions as control originating from clone names 13). CLuc luciferase was used as positive control for transfection and to measure inducibility. Cells are Tet-Off so they are growing in the presence of 2 μg doxycycline that is the homolog of tetracycline; G418 (400 μg/ml) was used to select and maintain the reporter gene carrying cells (for inducibility). Hygromycin (50 to 200 μg/ml) is used to select the transfected cells and maintain the cells carrying the transfected construct such as morpheus C5 and luciferase CLuc. The MEF cells are transfected with the respective constructs and kept under a continuous feeding and selection process. Tet-Off MEF cells were purchased from Clontech.

### Immunoprecipitation

Cells are transfected with either GFP or GFP-NPIP constructs. 24 hours after transfections, cells are lysed with 1 ml of RIPA buffer; 15 μl of this lysate is separated for western blot analysis. 60 μl of protein A beads (Amersham) are added to the lysate and incubated for 1 hour at 4°C with inversion. The slurry is spun down for 1 min at 4°C. The pre-cleared lysate is incubated with 2.5 μg of primary NPIP antibodies (N5 and N8) or preimmune N8. The antibody and lysate mixture is incubated for 1 hour at 4°C with inversion. 60 μl of 50% protein A beads are added on the antibody-lysate mixture and left at 4°C overnight. The tubes are spun down (1 min full speed) and the beads are washed three times with 1 ml lysis solution. After the beads are washed, 5X SDS loading buffer is added and boiled for 5 min before the SDS-PAGE analysis.

### Subcellular fractionation

Cells are grown on a 5 cm plate until they are 90% confluent (approx. 1 million cells). Cytoplasmic, membrane, nucleus and cytoskeleton (or precipitated proteins) are fractionated with Qproteome cell compartment kit (Qiagen) and run on a SDS-PAGE gel and detected by western blot. The cells are treated with 100 μg/ml pI:C (mimics viral infection) for 24 hours. Cells are scraped into media or PBS and precipitated by centrifugation at 300 g for 5 min at RT. The supernatant is discarded and the pellet is subjected to biochemical fractionation. At the end of this fractionation, the protein lysate is precipitated with acetone to be able to run them on a SDS PAGE gel. Precipitated lysate is stored at −20°C until SDS-PAGE analysis.

### Western blot analysis

Proteins were run on SDS-PAGE gel and transferred to nitrocellulose membrane by electroblotting. Ponceau-S (0.1% Ponceau-S (w/v) (Sigma), in 5% acetic acid) staining was used to define the location of the proteins on a nitrocellulose membrane. The membrane was blocked with 5% milk powder, 0.1% Tween 20, for 15 hours at 4°C. Antiserum/antibody was diluted in PBS, 5% milk powder, 0.1% Tween-20, and protein bands were visualized using the enhanced chemiluminescence (ECL) substrate (Millipore).

### Immunofluorescence

Appropriate cell lines grown on 22X22 mm coverslips in 6-well plates were induced, left uninduced with interferon γ, or transfected with NPIP constructs. After 24 hours, the medium was removed. The cells were washed with 2 ml of PBS and fixed with 2 ml of PBS/3%Paraformaldehyde for 20 min at room temperature. Cells were washed three times with PBS and washed with 2 ml PBS/0.1% Saponin incubated for 10 min at room temperature. The wash buffer was removed and immediately cells were blocked by adding PBS/0.1%Saponin/3% BSA and incubated for 1 hour at room temperature in 6-well plates. Coverslips were incubated with 100 μl of PBS/0.1%Saponin/3% BSA, which contains the appropriate antibody dilution on Parafilm in a humidified chamber for 1 hour at room temperature or overnight at 4°C. Coverslips were put in to the 6-well plates and washed with 3X 5 ml of PBS/0.1% Saponin. Coverslips were incubated with 100 μl of PBS/0.1% Saponin/3% BSA, which contains the appropriate secondary antibody dilution or DAPI (1:1000) on Parafilm in a humidified chamber for 30 min at room temperature in the dark. Coverslips were put into the 6-well plates and washed with 3X 5 ml of PBS/0.1% Saponin. Finally, coverslips were put on to the slide with 20 μl of ProLong Gold antifade reagent (Molecular Probes). After overnight incubation, cells were observed with a Leica (DM4000B) fluorescence microscope equipped with a CCD camera (DFC350FX) using the Leica software (LeicaApplicationSuite, 2.1.0 build 4316).

## ACKNOWLEDGMENTS

This work was supported, in part, by the National Institutes of Health [grant numbers GM058815, HG002385 to EEE]; by TUBITAK-1001 [grant number 112T421 to CB]; and by the Intramural Research Program of the National Human Genome Research Institute, National Institutes of Health to NISC and JCM. We would like to thank Cezmi A. Akdis for his technical support and Tonia Brown for feedback on the manuscript. EEE is an investigator of the Howard Hughes Medical Institute.

## SUPPLEMENTARY FIGURES

**Figure S1.**
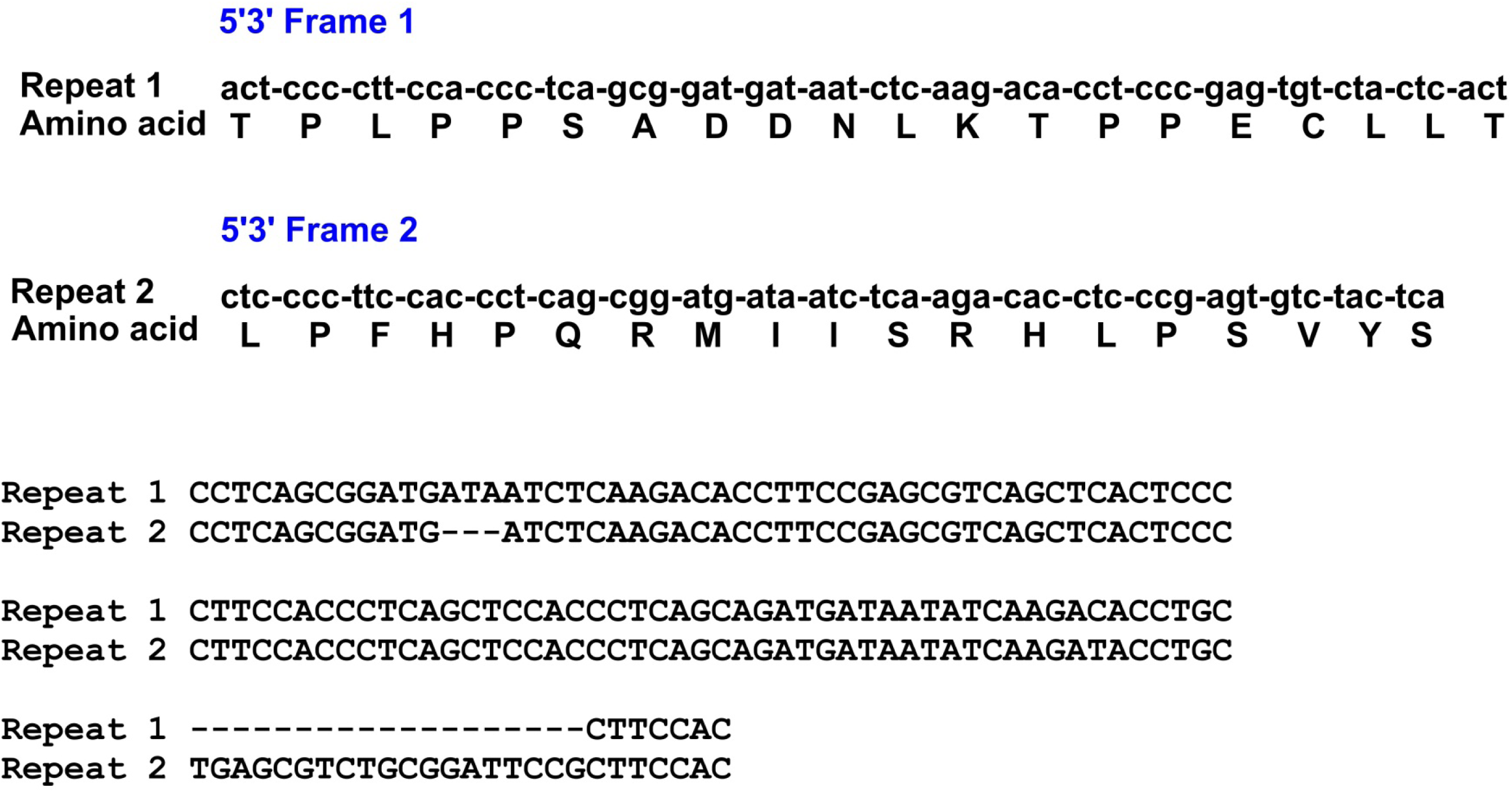
Frameshift mutation in the repeat region of NPIPA and NPIPB

**Figure S2.**
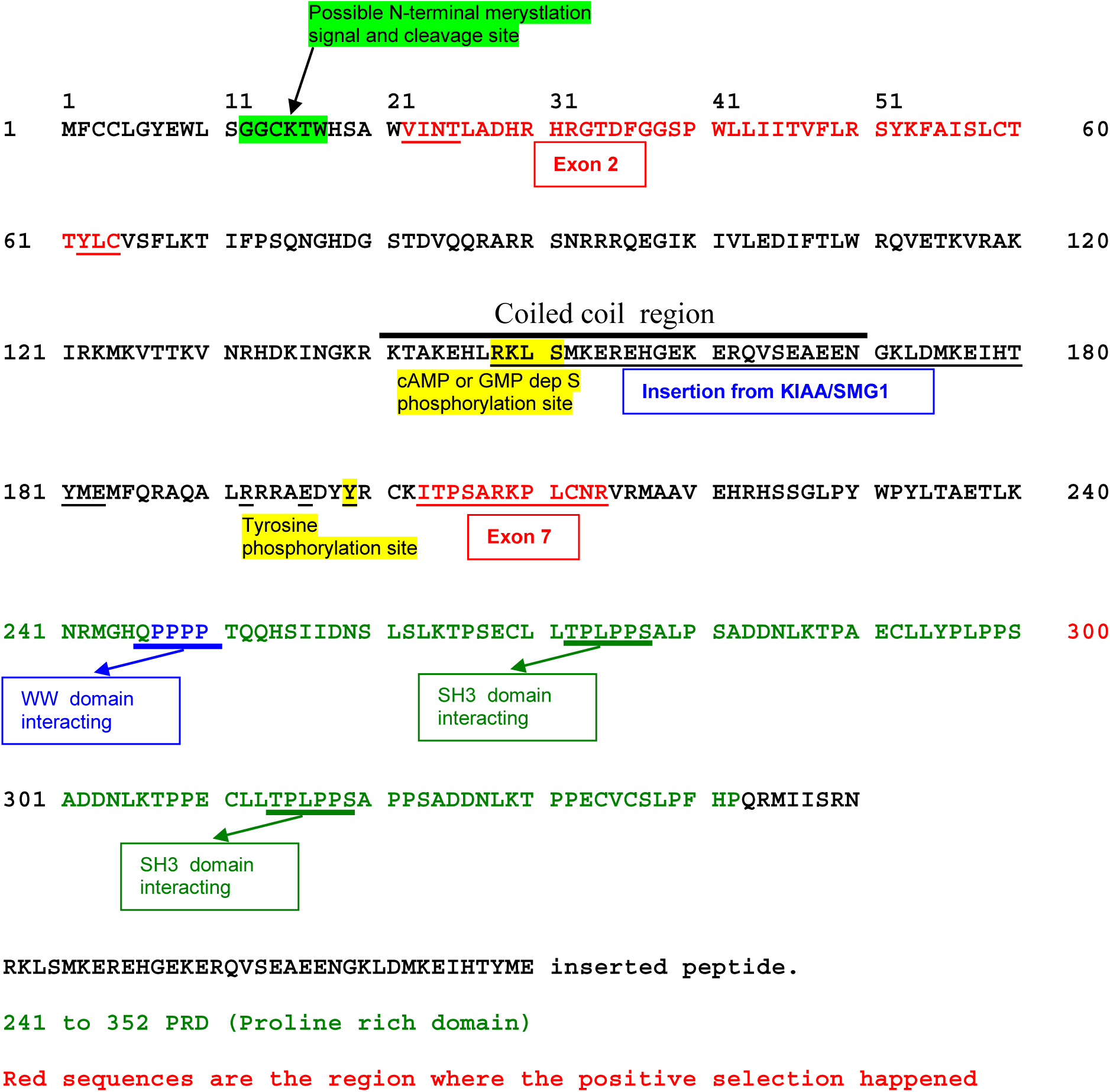
Figure S2. Morpheus protein domains and phosphorylation sites

**Figure S3.**
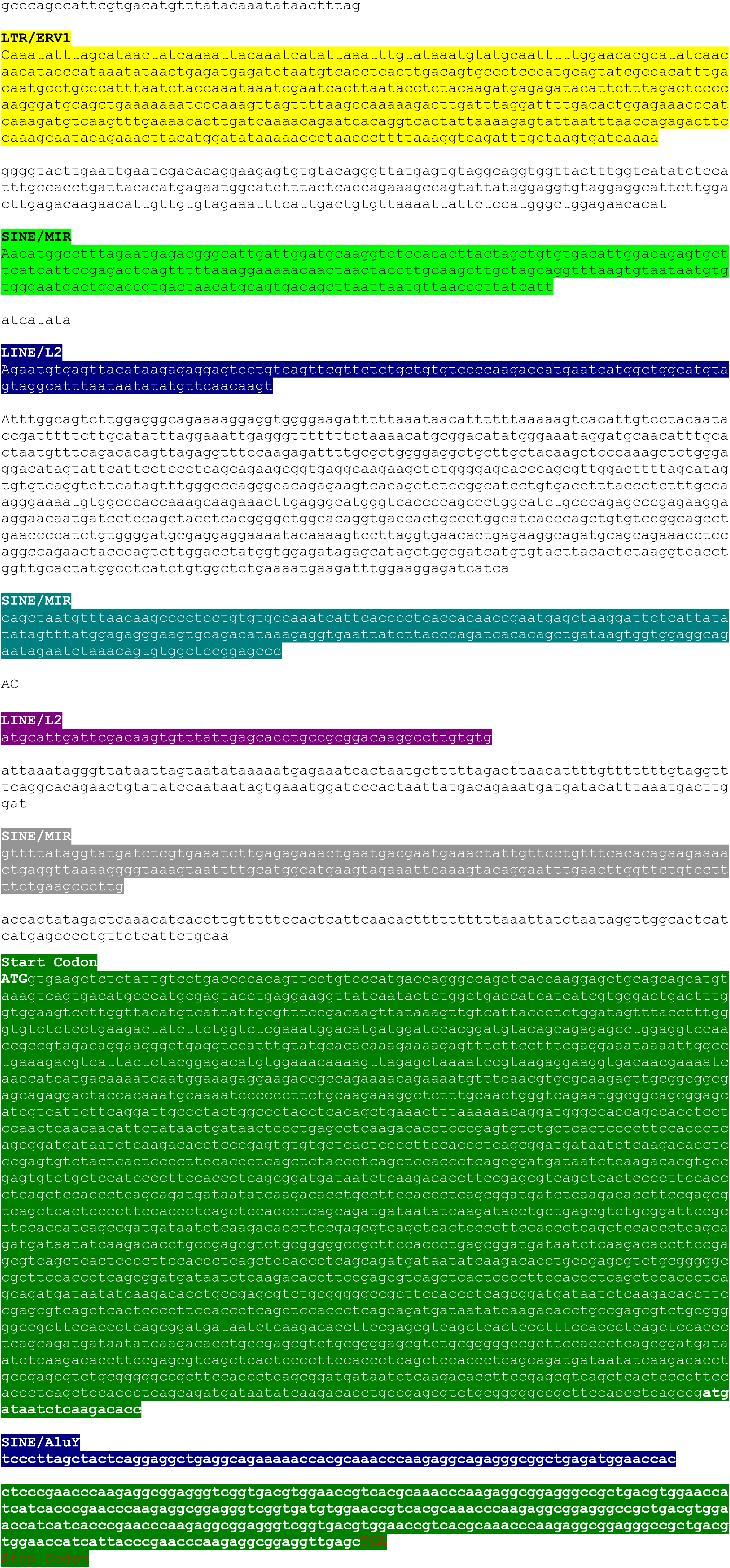
5’UTR repeat content.

**Figure S4.**
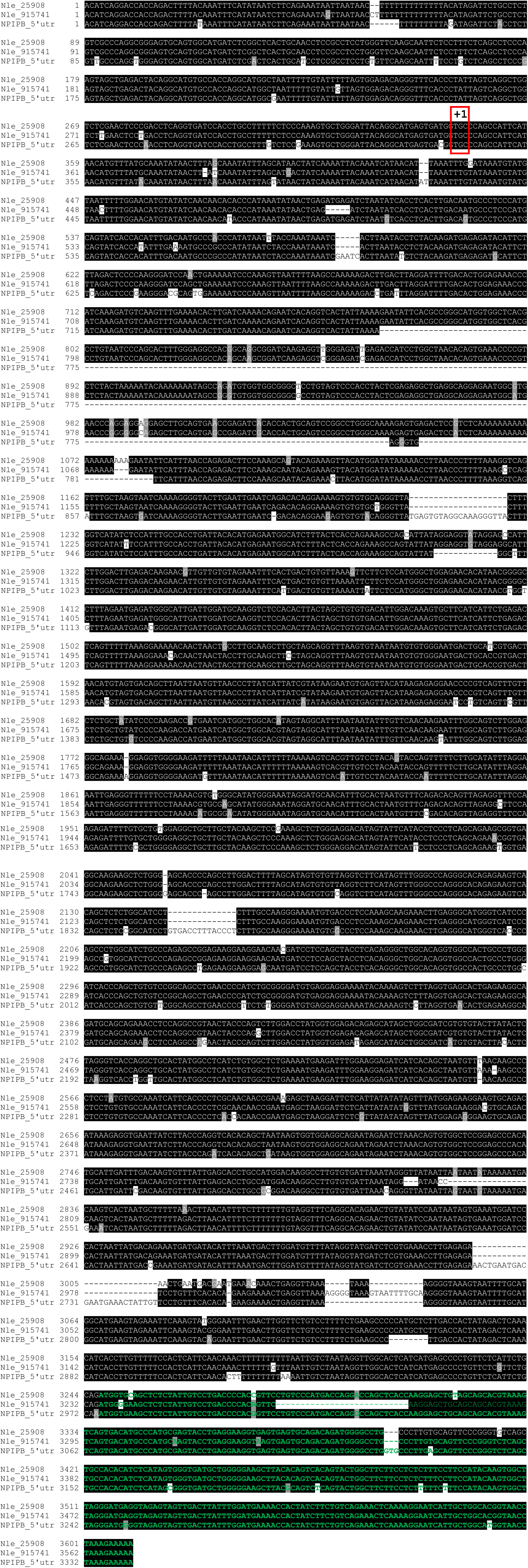
Alignment of the 5’UTR including the promoter side

**Figure S5.**
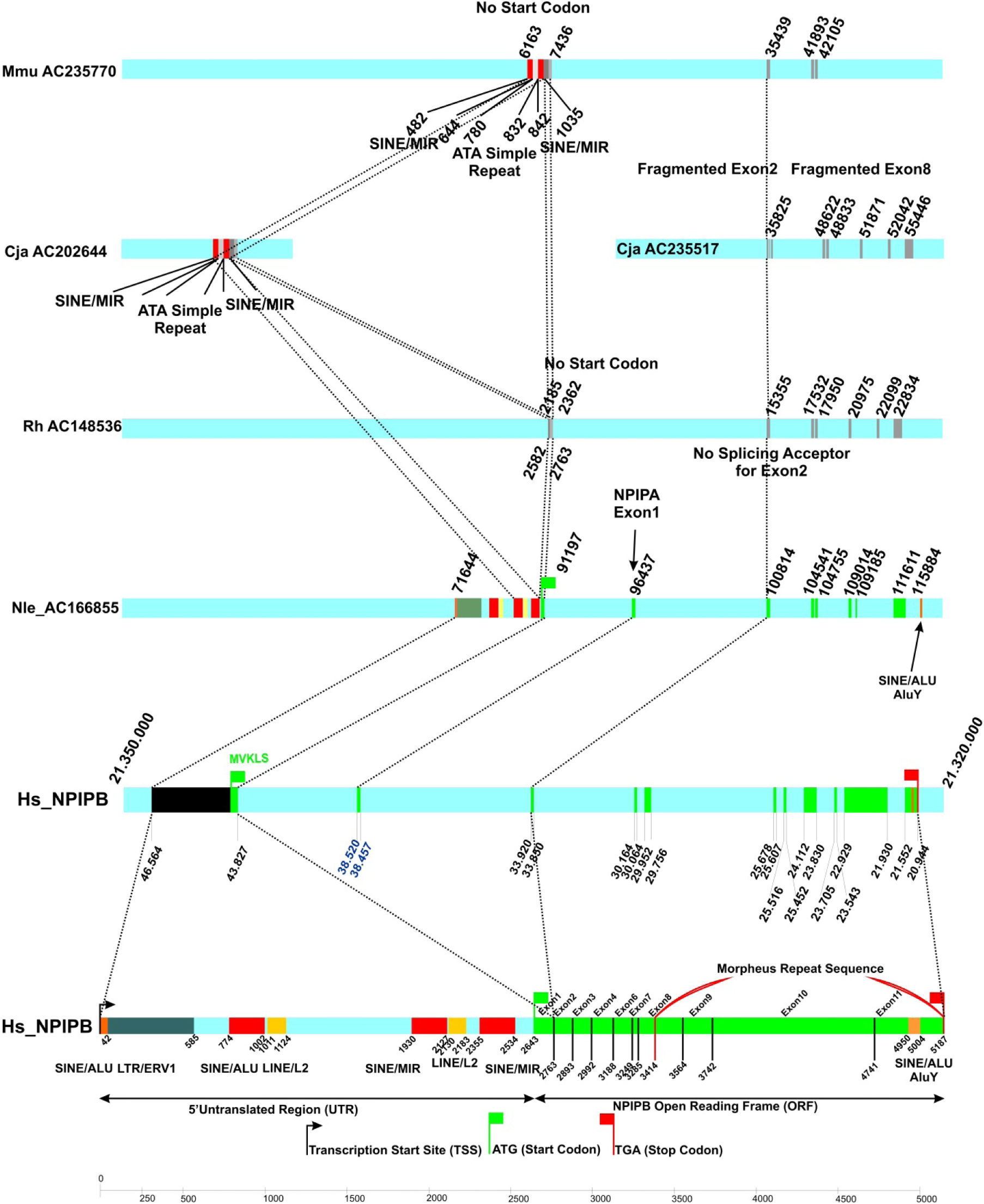
Comparative genomics analysis of prosimian, marmoset, Rhesus macaque, gibbon and human. The figure indicates the different gene structure of NPIP from Mmu *(Microcebus murinus,* AC235770), Cja (*Calithrix jacchus,* AC202644, AC235517), Rh, (*Macaca mulatta,* AC148536), (*Nomascus leucogeny,* AC166855) and human (*Homo sapiens*).

**Figure S6.**
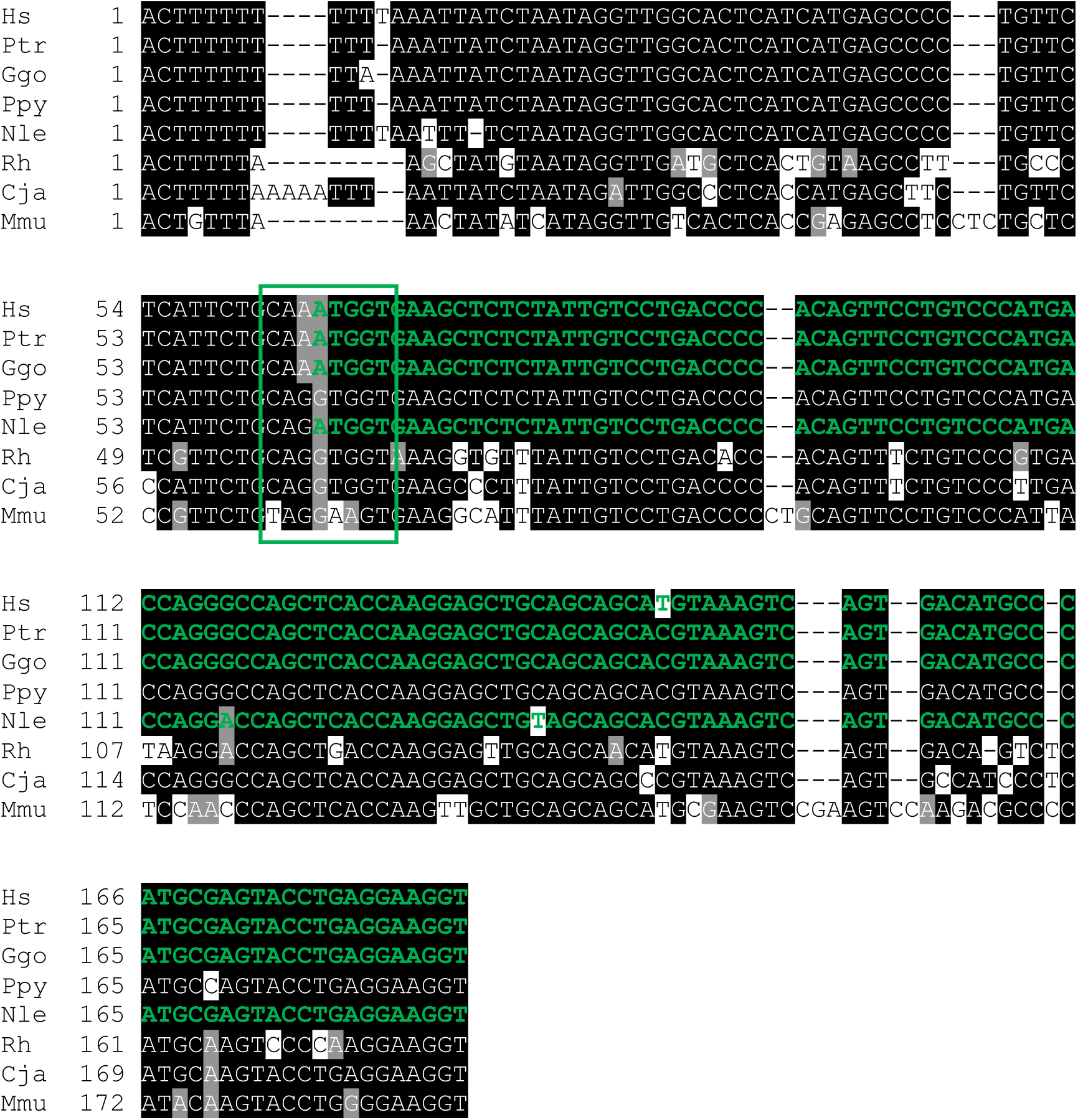
Alignment of 1^st^ exon of NPIPB gene, including the partial 5’UTR. Sequence that encodes the proper morpheus protein is bolded and colored in green. The green box indicates the position of start codon.

**Figure S7.**
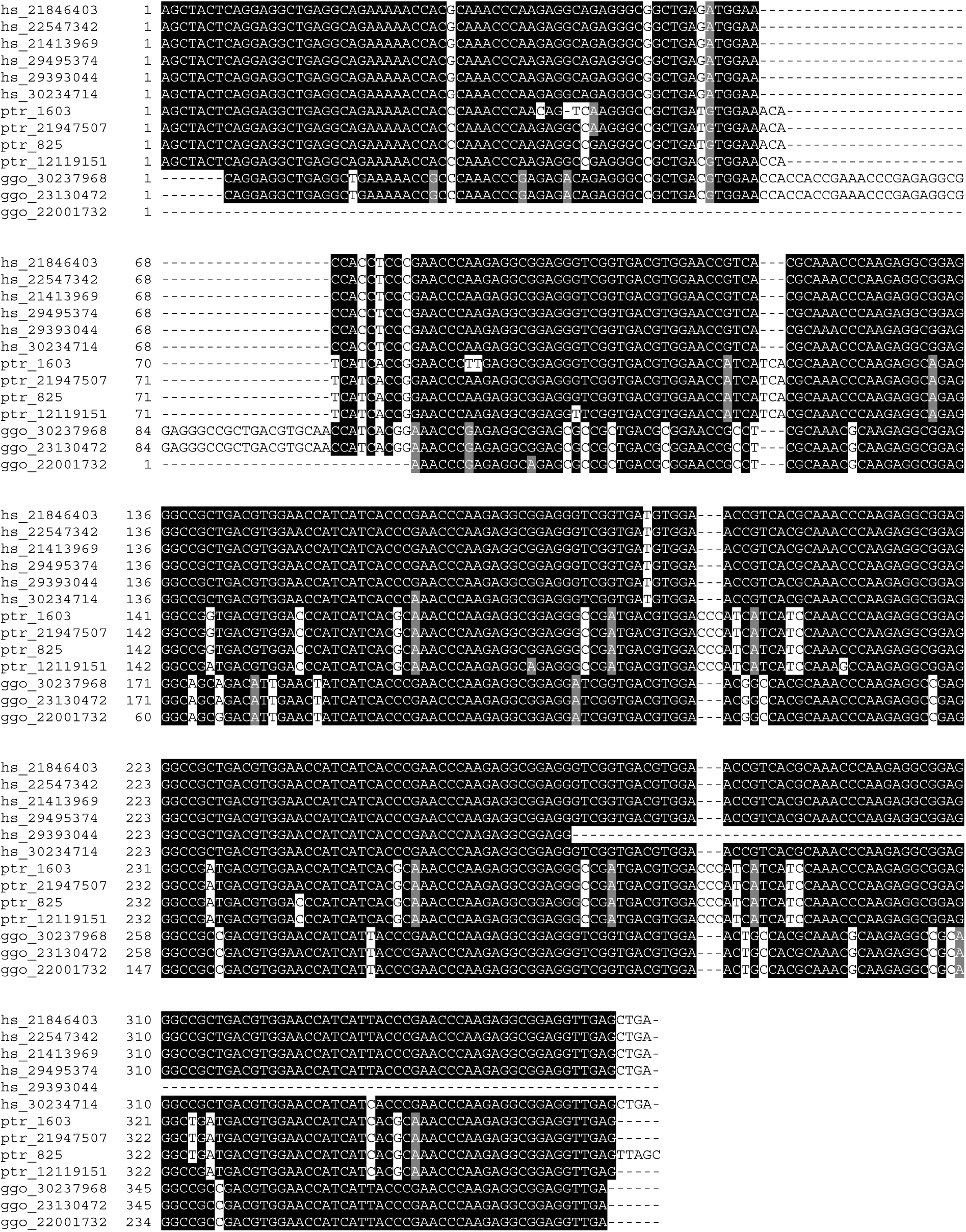
Alignment of repeat 3’UTR including the AluY element

**Figure S8.**
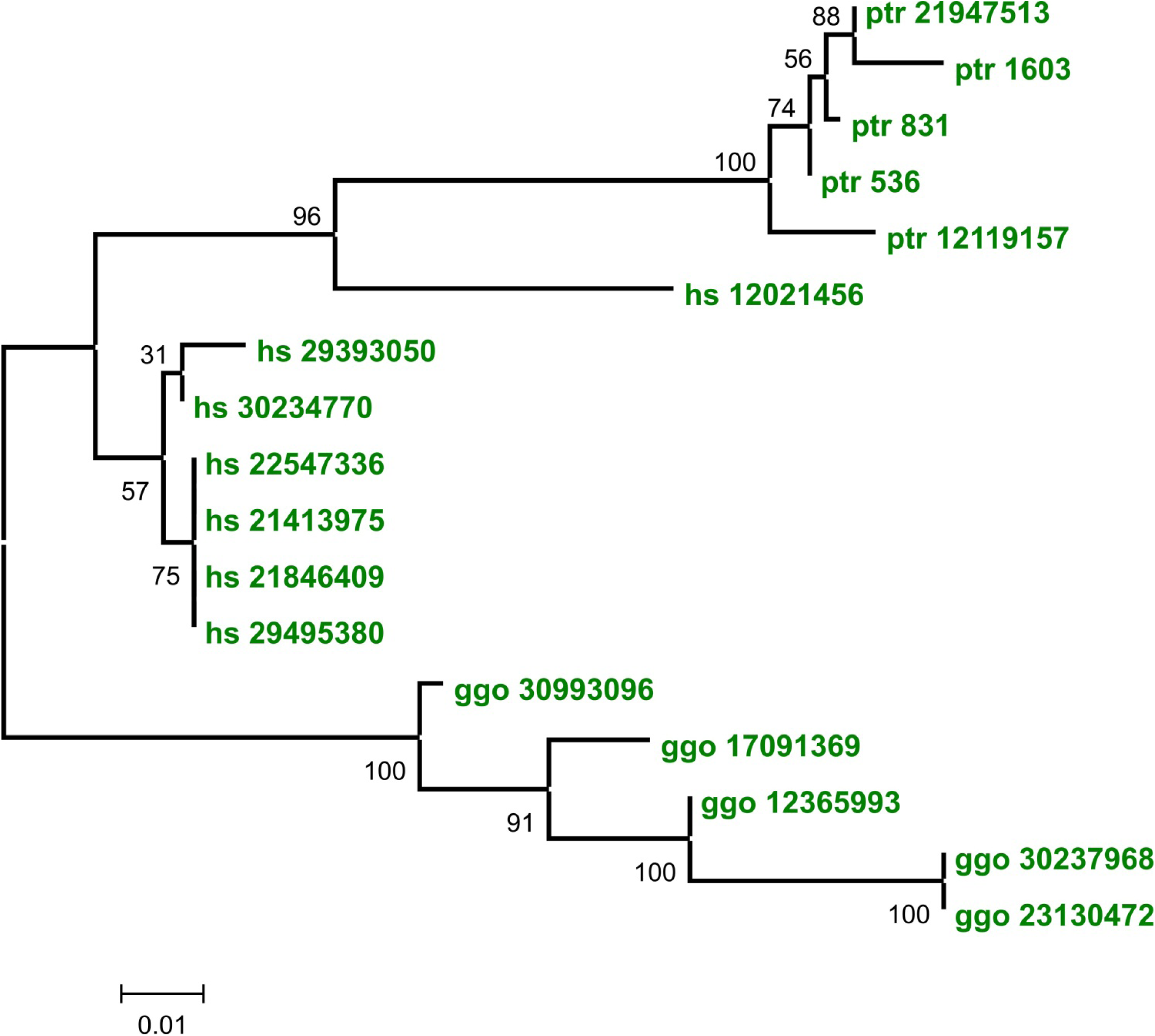
Phylogeny of repeat 3’UTR with AluY element. The evolutionary history was inferred using the neighbor-joining method. The optimal tree with the sum of branch length = 0.31266271 is shown. The percentage of replicate trees in which the associated taxa clustered together in the bootstrap test (500 replicates) are shown next to the branches (19). The tree is drawn to scale, with branch lengths in the same units as those of the evolutionary distances used to infer the phylogenetic tree. Codon positions included were 1st+2nd+3rd+Noncoding. All positions containing alignment gaps and missing data were eliminated only in pairwise sequence comparisons (pairwise deletion option). There were a total of 413 positions in the final dataset. Phylogenetic analyses were conducted in MEGA4 (21).

**Figure S9.**
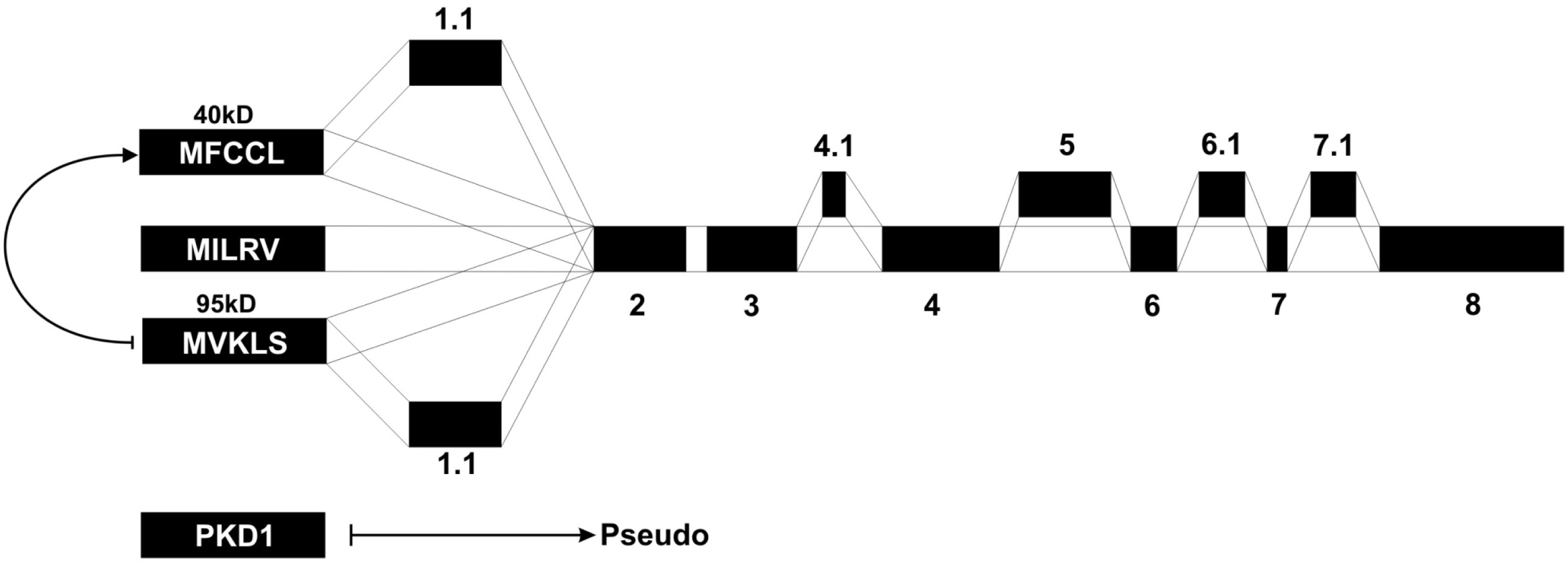
Alternative splicing for morpheus gene family. Splicing structure for the morpheus gene family is illustrated. All possible alternative exon formations are shown with the connected lines. Morpheus-PKD1 fused gene always produces a pseudogene formation. It is also possible that long version 95 kDa splices into short splice version.

**Figure S10.**
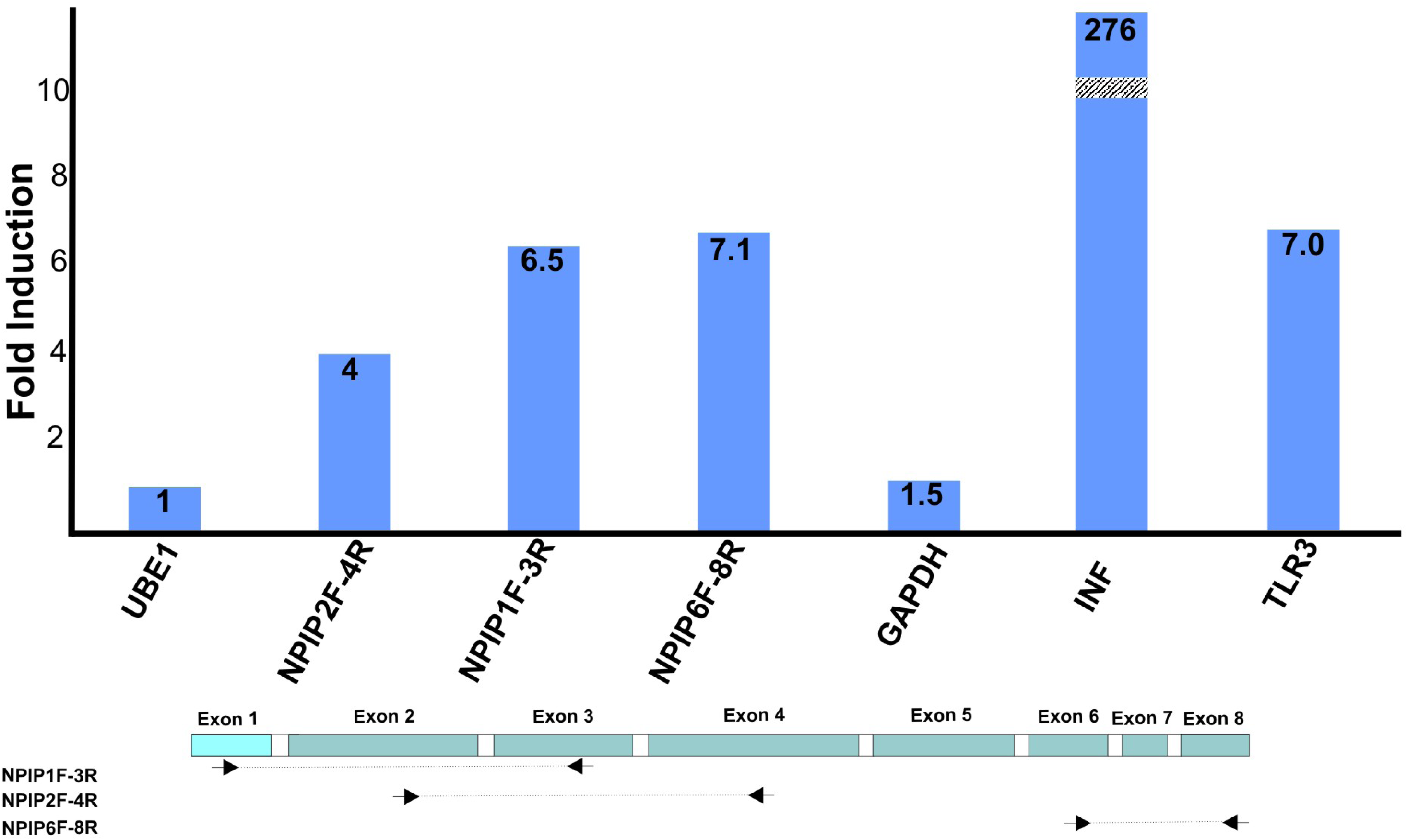
Induction of morpheus gene family by pI:C transfection. Real-time PCR analysis to show fold induction of morpheus gene family between pI:C induced and uninduced HeLa cells. HeLa cells are transfected with 1μg pI:C for 6 hours and harvested by scraping to media followed by ctg. 300 g at RT for 5 min. Real-time PCR is performed on cDNA prepared from the total RNA (Qiagen RNeasy kit) of induced and uninduced HeLa cells. UBE1 and GAPDH are used for control. Interferon (INF) and toll-like receptor 3 (TLR3) genes are used as positive control. The primers for real-time PCR were designed for exon 6 and 8.

**Figure S11.**
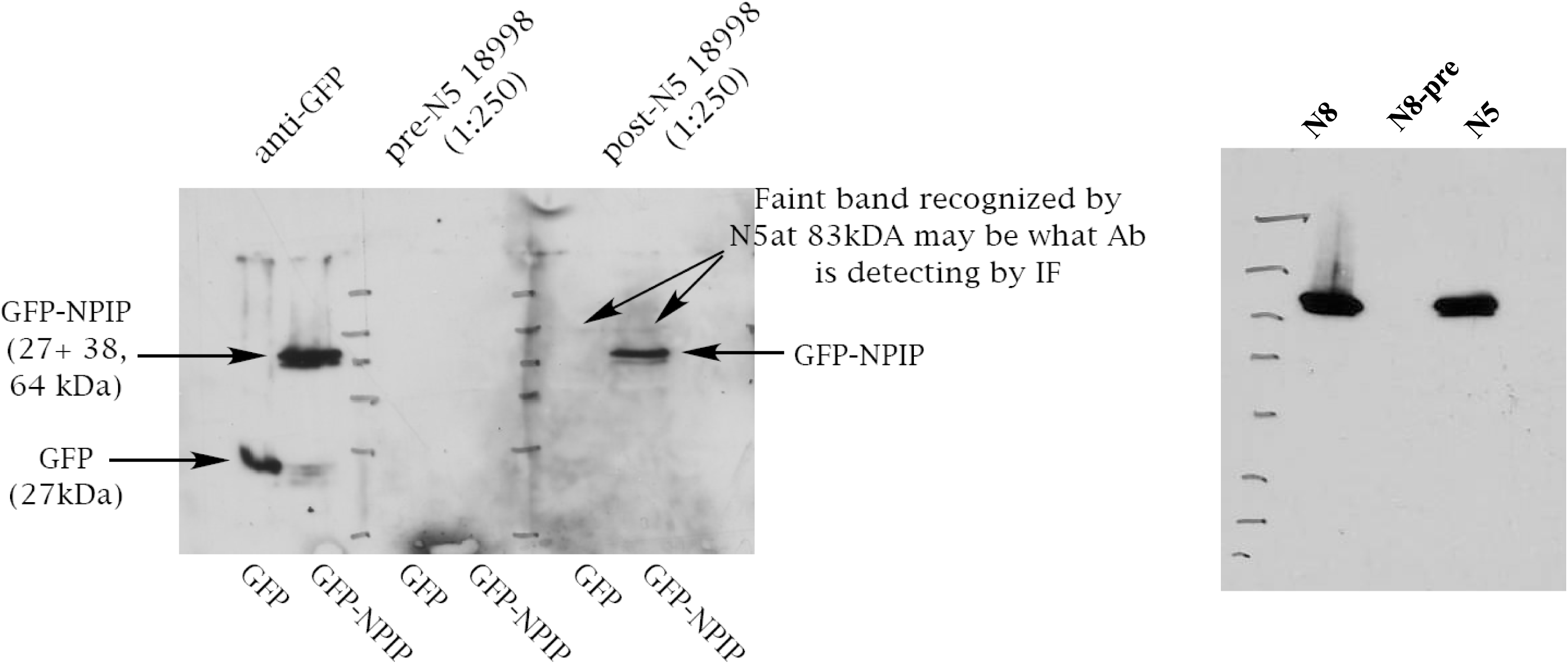
N5 and N8 antibodies specifically recognize the transfected GFP-NPIP fusion protein. N5 and N8 antibodies specifically recognize the recombinant GFP-NPIP protein but not the pre-immune serum. Recombinant GFP-NPIP fusion protein that is expressed in HeLa cells is detected by N5 (left) and N8 (right) antibodies and not pre-immune antibody. Cells are transfected either with GFP-NPIP fusion construct or GFP alone and the recombinant GFP-NPIP fusion protein is immunoprecipitated with N5, N8, pre-immune serum for N5 and pre-immune serum for N8 (Material and Methods). The SDS-PAGE gel is blotted with anti-GFP (monoclonal) antibody.

**Figure S12.**
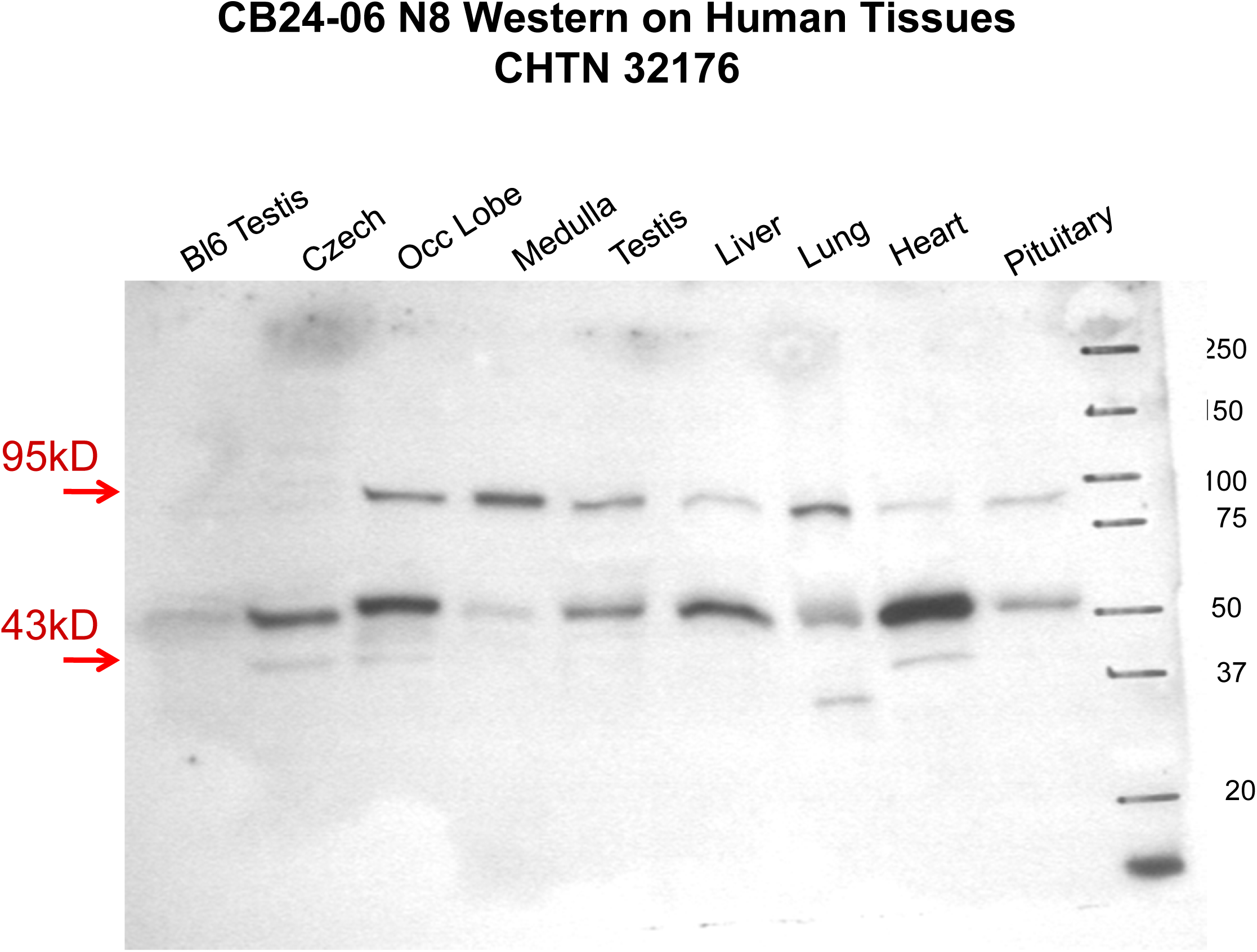
Western blot data showing human tissues compared to Bl6 mouse tissues. About 50 mg of each tissue (occipital lobe, medulla, testis, liver, lung, heart, and pituitary) and human lymphocyte cell line-Czech lysate (CB23-06) were harvested and homogenized in RIPA buffer by tissue homogenizer (Omni International, TH). The amount of the lysate was measured by BCA, Protein Assay Kit (Pierce). Each well is loaded with 50 μg of protein lysate. The SDS-PAGE gel was blotted on nitrocellulose membrane and stained with the respective antibody (N8) by using Pierce western blot detection kit (West Femto, HRP and ECL reagents).

**Figure S13.**
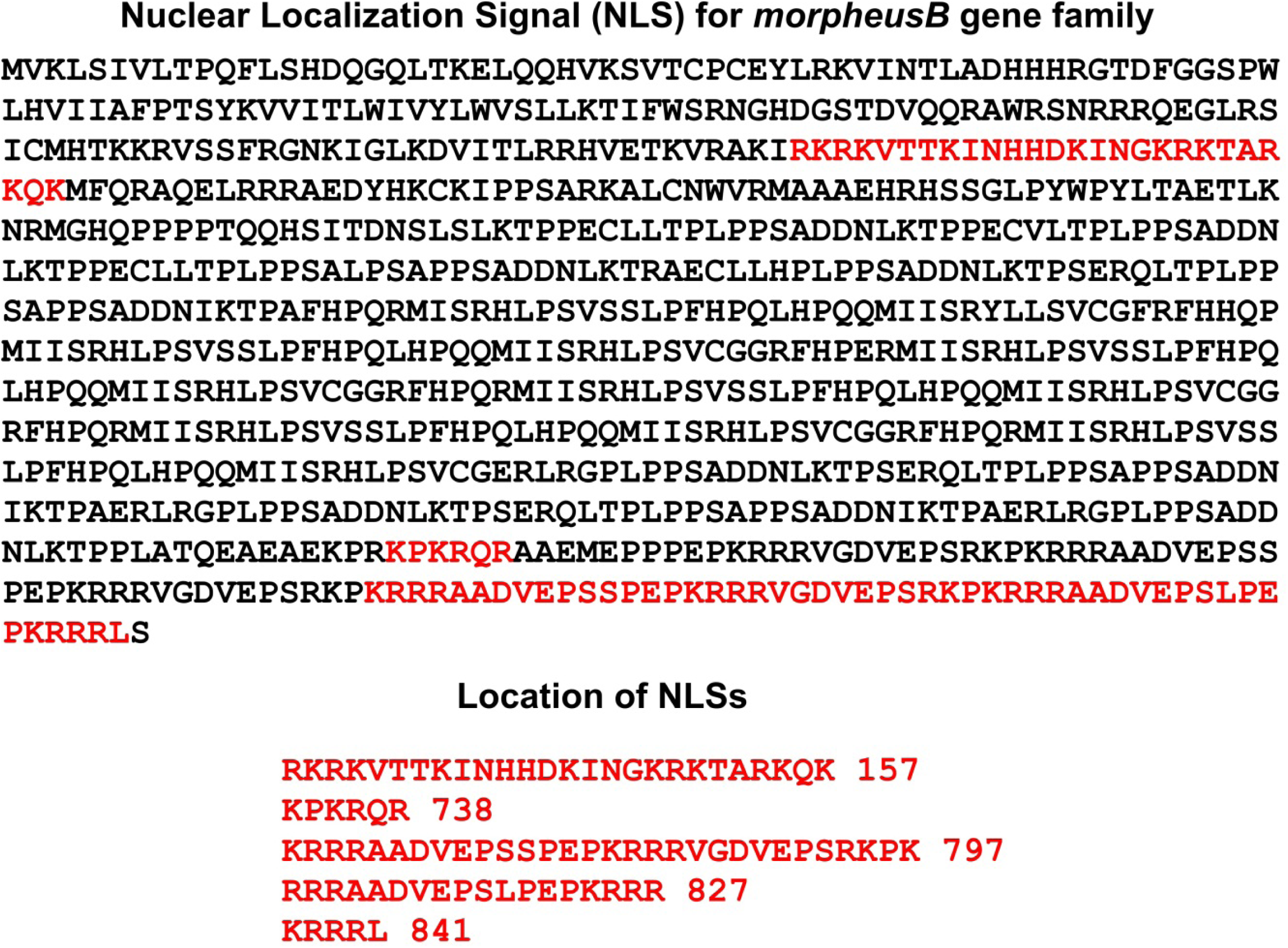
NLS signal for morpheus proteins. Putative NLS signal for NPIPB protein is predicted by using the web browser “PredictProtein”.

**Figure S14.**
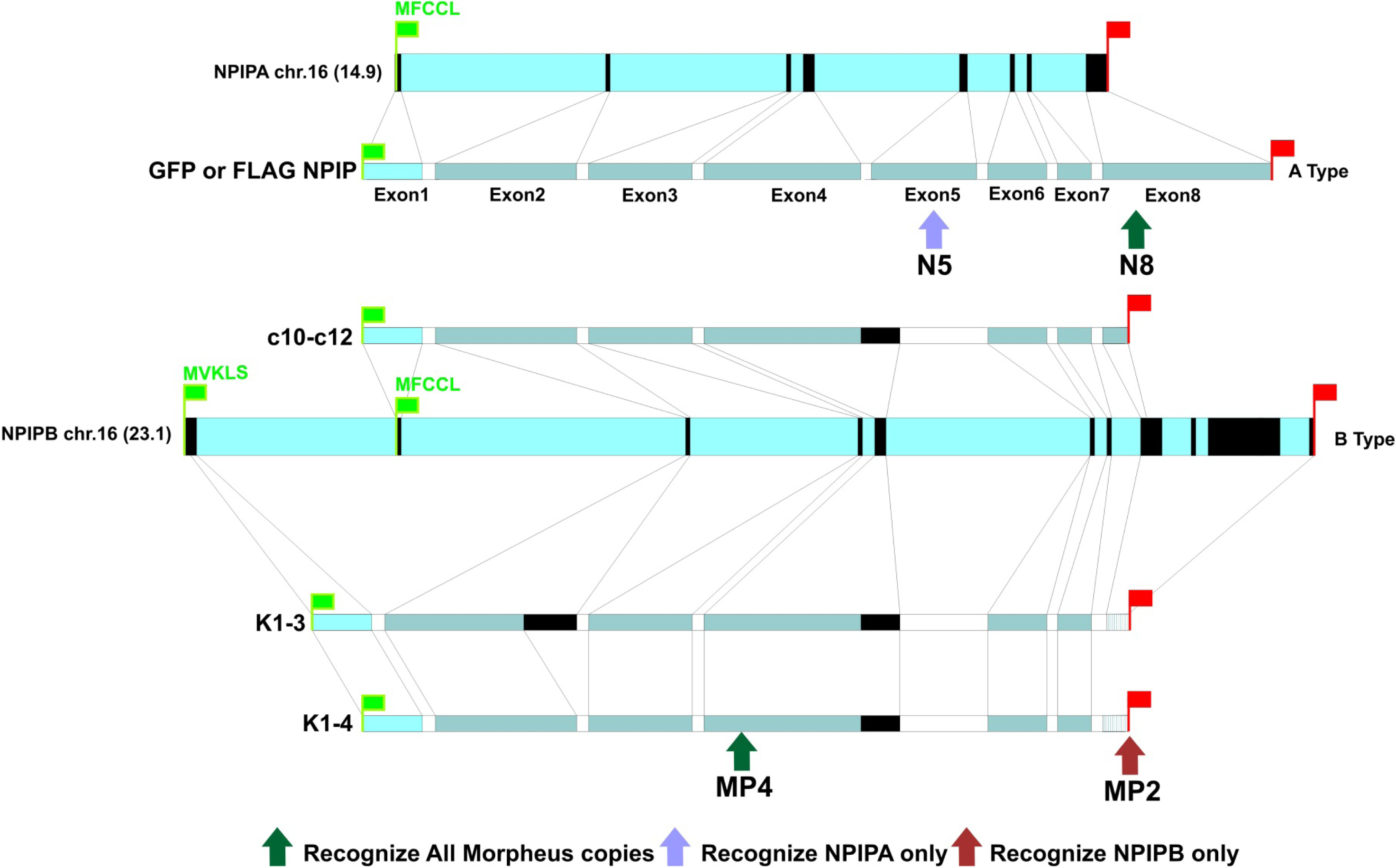
The comparative gene structure for GFP Flag-NPIP, c12, K1-3 and K1-4 constructs on *NPIPA* and *NPIPB* gene structure. The figure shows the comparative gene structure of GFP Flag-NPIP and c12 constructs depicted on long *NPIPB* (95 kDa) and short *NPIPA* (40 kDa) (NPIP) (see Figure 1). The green and red flags indicate the position of start and stop codons, respectively. The GFP Flag-NPIP gene is different from the c12 construct by having Exon 5 and additional sequence at C-terminal region. NPIPB, K1-3 and K1-4 constructs are only different with the long exon2 and C terminal repeat region.

**Figure S15.**
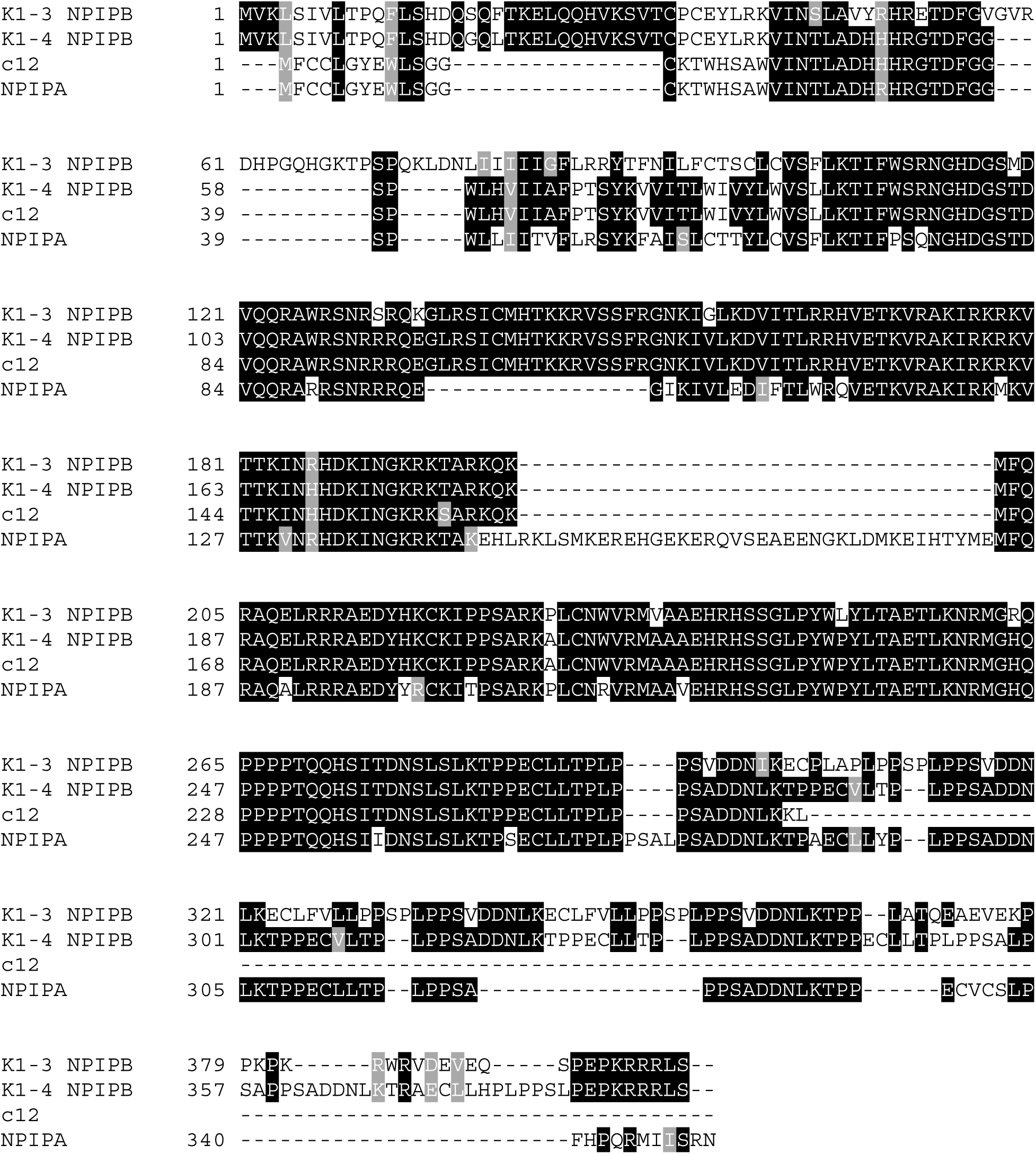
Alignment of c12, NPIP, K1-3 (NPIPB) and K1-4 (NPIPB)

**Figure S16.**
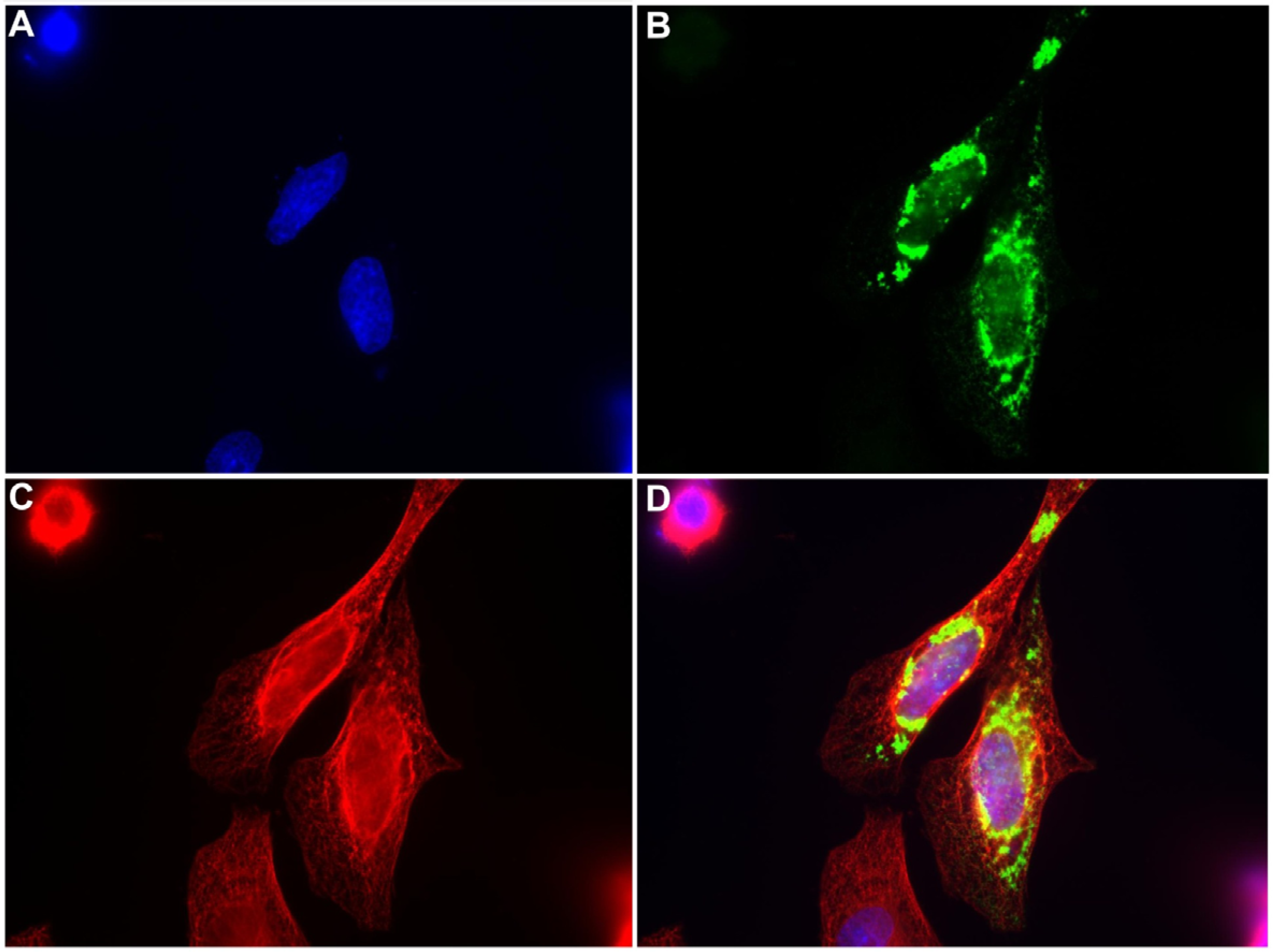
Immunofluorescence analysis of “c12” gene in HeLa cells. HeLa cells are transiently transfected by using the FuGENE HD transfection reagents (Roche) with c12 for 24h and stained with 1–100 dilution of N8 antibody (B) and 1–1000 dilution of anti-tubulin antibody (C). Primary antibody staining is performed for overnight at 4°C. Nucleus (A) is shown by DAPI staining. Merged figure (D) shows the co-localization between N8 (B) and tubulin antibody (C) staining. The majority of the HeLa cells are showing specific staining within the nucleus.

**Figure S17.**
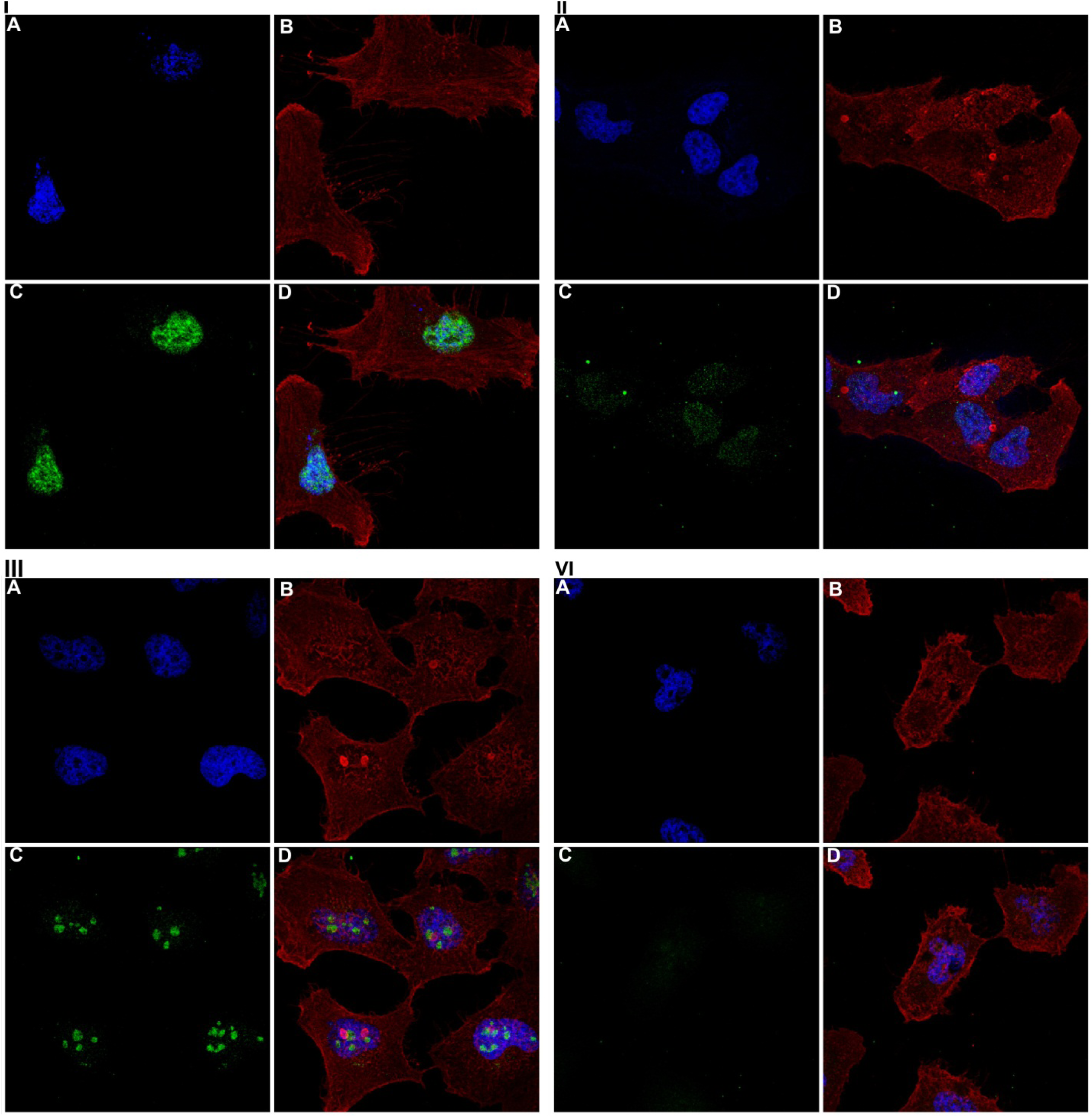
Endogenous level of expression of morpheus B protein in HeLa cells. Immunofluorescence analysis is shown of *NPIPB* in HeLa cells. HeLa cells are stained with 1–500 dilution of MP4 (I-C) and preimmune antibody (II-C), MP2 (III-C) and preimmune antibody (IV-C), and a 1–1000 dilution of anti-beta-actin antibody (Cell Signaling) (B). The nucleus (A) is visualized by DAPI staining. The merged figure (D) shows the co-localization between MP4 (C) and beta actin antibody (B) staining. The majority of the HeLa cells show specific staining within the nucleus.

**Figure S18.**
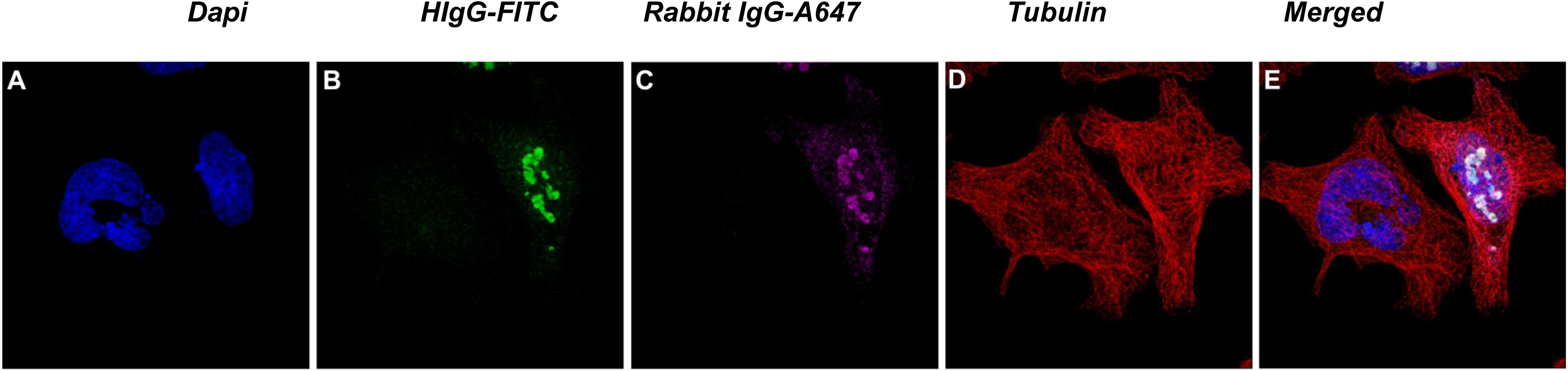
Detection of human IgG for morpheus protein. HeLa cells are transiently transfected by using the FuGENE HD transfection reagents (Roche) with K1-3 for 24h and stained with 1–1000 dilution of proteinA isolated human IgG from healthy individual, 1–500 dilution of MP4 antibody (C) and 1–1000 dilution of anti-tubulin antibody (sigma) (D). Primary antibody staining is performed for overnight at 4°C. Human IgG fraction is detected by using GoatF(ab)2 antihuman IgG (γ)FITC Conjugate (Invitrogen), Anti-rabbitIgGAlexa 647 Conjugate. Nucleus (A) is shown by DAPI staining. Merged figure (E) shows the co-localization between Human IgGMP4 (C), and beta actin antibody (B) staining.

**Supplementary Table 1.** Morpheus domain search SMART results

| Supplementary Table 1 Morpheus Domain SMART results |  |  |  |  |
| --- | --- | --- | --- | --- |
| **Domains for NPIPB** |  |  |  |  |
| Name | Domain information | Begin | End | E-value |
| HALZ | Homeobox associated leucine zipper | 29 | 77 | 7.29e+02 |
| IRF | Interferon regulatory factor | 68 | 159 | 2.49e+03 |
| Hr1 | Rho effector or protein kinase C-related kinase homology region 1 homologues | 138 | 205 | 8.05e+02 |
| IENR2 | Intron-encoded nuclease repeat 2 | 145 | 161 | 2.19e+03 |
| Ubox | Modified RING finger domain | 443 | 496 | 1.79e+03 |
| Amelogenin | cell adhesion proteins, play a role in the biomineralisation of teeth. | 517 | 669 | 2.41e+04 |
| PLCXc | Phospholipase C, catalytic domain (part); domain X | 547 | 644 | 1.83e+03 |
| Ad_cyc_gAlp | Adenylate cyclase G-alpha binding domain | 700 | 733 | 1.36e+05 |
| **Domains for the NPIPA** |  |  |  |  |
| IL2 | IL2 family | 52 | 229 | 2.12e+03 |
| HSA | domain in helicases and associated with SANT domains | 86 | 152 | 1.60e+03 |
| BRLZ | Basic region leucine zipper | 81 | 139 | 3.19e+02 |
| Hr1 | Rho effector or protein kinase C-related kinase homology region 1 homologues | 97 | 163 | 1.52e+03 |
| CRA | CT11-RanBPM | 138 | 240 | 8.40e+06 |
| Zf-AD | Zinc finger associated domain | 173 | 195 | 6.85e+04 |
| DM6 | Cysteine-rich domain currently specific to Drosophila. | 183 | 306 | 3.03e+03 |
| Amelogenin | cell adhesion proteins, play a role in the biomineralisation of teeth. | 222 | 336 | 2.41e+04 |

**Supplementary Table 2.**
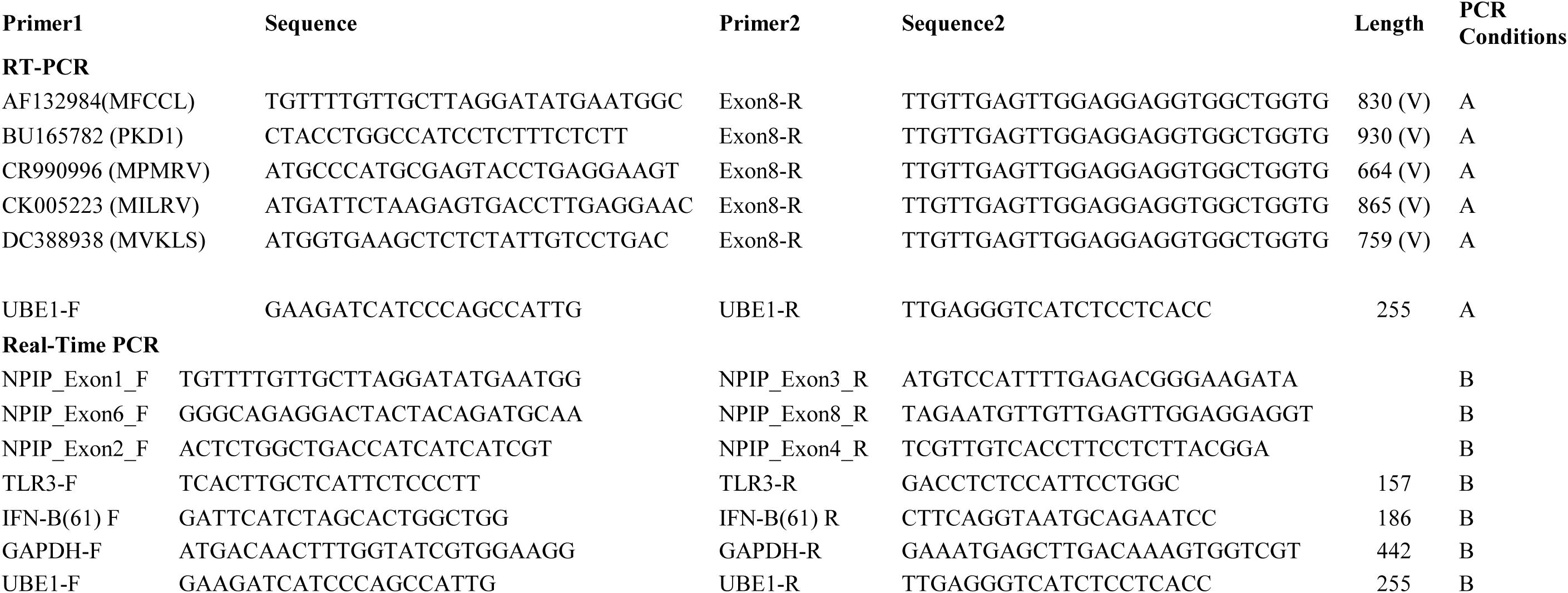
Primer table for morpheus genes

